# Mutational bias in spermatogonia impacts the anatomy of regulatory sites in the human genome

**DOI:** 10.1101/2021.06.10.447556

**Authors:** Vera B. Kaiser, Lana Talmane, Yatendra Kumar, Fiona Semple, Marie MacLennan, Deciphering Developmental Disorders Study, David R. FitzPatrick, Martin S. Taylor, Colin A. Semple

## Abstract

Mutation in the germline is the ultimate source of genetic variation, but little is known about the influence of germline chromatin structure on mutational processes. Using ATAC-seq, we profile the open chromatin landscape of human spermatogonia, the most proliferative cell-type of the germline, identifying transcription factor binding sites (TFBSs) and PRDM9-binding sites, a subset of which will initiate meiotic recombination. We observe an increase in rare structural variant (SV) breakpoints at PRDM9-bound sites, implicating meiotic recombination in the generation of structural variation. Many germline TFBSs, such as NRF, are also associated with increased rates of SV breakpoints, apparently independent of recombination. Singleton short insertions (>=5 bp) are highly enriched at TFBSs, particularly at sites bound by testis active TFs, and their rates correlate with those of structural variant breakpoints. Short insertions often duplicate the TFBS motif, leading to clustering of motif sites near regulatory regions in this male-driven evolutionary process. Increased mutation loads at germline TFBSs disproportionately affect neural enhancers with activity in spermatogonia, potentially altering neurodevelopmental regulatory architecture. Local chromatin structure in spermatogonia is thus pervasive in shaping both evolution and disease.

## Introduction

Mutation is the ultimate source of genetic variation, and inherited variation must invariably arise in the germline. It is well established from cross-species comparisons that the rate of nucleotide substitution mutations fluctuates at the multi-megabase (>10^6 bp) scale across the genome (Wolfe et al. 1989; Hodgkinson and Eyre-Walker 2011), with early replicating regions subject to reduced rates of mutation. These patterns similarly manifest in the rate of human single nucleotide polymorphisms (SNPs) (Stamatoyannopoulos et al. 2009). Germline structural variation in the human genome is also associated with replication timing, such that copy number variants (CNVs) emerging from homologous recombination-based mechanisms are enriched in early replicating regions, while CNVs arising from non-homologous mechanisms are enriched in late replicating regions (Koren et al. 2012). Local chromatin structure also influences the mutation rate. However, finer-scale variation (<1Mb) in the germline mutation rate has so far only been related to genomic features derived from somatic cells (Gonzalez-Perez et al. 2019) because human germline-derived measures of chromatin structure have only recently become available (Guo et al. 2017; Guo et al. 2018). Transcription factor binding sites (TFBSs) are particularly prone to point mutations in cancer (Kaiser et al. 2016), probably due to interference between TF binding and the replication and repair machinery (Reijns et al. 2015; Sabarinathan et al. 2016; Afek et al. 2020), but the mutational consequences of binding at these sites in the germline is unknown.

During meiosis, homologous recombination may introduce short mutations or render genomic regions prone to rearrangements (Pratto et al. 2014; Halldorsson et al. 2019). A key player in this process is PRDM9, which binds its cognate sequence motif and directs double-strand break (DSB) formation in meiotic prophase (Baudat et al. 2010; Myers et al. 2010). In humans, PRDM9 binding site occupancy has only been directly assayed in a somatic cell line (Altemose et al. 2017), whereas indirect measures of PRDM9 activity include a proxy for DSBs (DMC1-bound single stranded DNA (ssDNA)) in testis (Pratto et al. 2014), and population genetic based measures of recombination hotspots (HSs) (Myers et al. 2005; The 1000 Genomes Project Consortium et al. 2015). The method ATAC-Seq (Buenrostro et al. 2013) reports chromatin accessibility and provides a snapshot of all active regulatory regions and occupied binding sites in a given tissue. In particular, ATAC-Seq footprinting (Sherwood et al. 2014; Li et al. 2019), when applied to spermatogonia, has the potential to reveal the binding of hundreds of TFs, as well as PRDM9, in the male germline. In addition, large human genome sequencing projects can be used to reveal patterns of mutation rates, by focussing on extremely rare variants (Messer 2009; Carlson et al. 2018; Li and Luscombe 2020). Making use of such variant datasets as well as novel ATAC-Seq data in spermatogonia, we study the mutational landscape at transcription factor binding sites (TFBSs) in accessible human spermatogonial chromatin.

## Results

### Spermatogonial regulatory regions are enriched for rare deletion breakpoints

ATAC-Seq (Buenrostro et al. 2013) reports local chromatin accessibility and provides a snapshot of active regulatory regions and genomic regions occupied by DNA-binding proteins in a given tissue. We used ATAC-Seq to identify open chromatin sites in FGFR3-positive spermatogonial cells isolated from dissociated human testicular samples. FGFR3 is most highly expressed in self-renewing spermatogonial stem cells, with low expression also being detected in early differentiating spermatogonia (Guo et al. 2018; Sohni et al. 2019); its expression thus overlaps with the onset of PRDM9 expression in pre-meiotic spermatogonia (Human Protein Atlas: https://www.proteinatlas.org/ENSG00000164256-PRDM9/celltype/testis and https://www.proteinatlas.org/ENSG00000068078-FGFR3/celltype/testis) (Guo et al. 2018). Open chromatin in FGFR3-positive cells was identified using standard peak detection analysis (Methods) and multiple metrics (Supplemental Fig. S1) indicated high data quality (Yan et al. 2020). Hierarchical clustering (Ramirez et al. 2016) showed that this novel spermatogonial ATAC-Seq dataset displays a genome-wide distribution of peaks consistent with other spermatogonial derived data, and is distinct from ES cell and somatic tissue datasets (Supplemental Fig. S2).

Next, we assessed the enrichments of different classes of sequence variants at spermatogonial active sites, including singleton SV breakpoint frequencies as a proxy for the mutation rate of such variants. We made use of ultra-rare genomic variants from a variety of human sequencing studies: the Deciphering Developmental Disorders (DDD) study (Deciphering Developmental Disorders Study 2015; Mcrae et al. 2017) of severe and undiagnosed developmental disorders (https://www.ddduk.org/), a large collection of variants from an aggregated database (gnomAD; http://gnomad.broadinstitute.org/), and *de novo* variants from trio sequencing studies (http://denovo-db.gs.washington.edu/, https://research.mss.ng/, An et al. (2018)). Based on the DDD dataset - a combination of high-density arrayCGH and exome sequencing (Deciphering Developmental Disorders Study 2015) - we identified 6,704 singleton deletion variants among 9,625 DDD probands (carrier frequency of ∼ 0.002) (Supplemental Table S1). Permutation analysis demonstrates that DDD singleton breakpoints are enriched at spermatogonial ATAC-Seq sites, their overlap being > 4-times the expected genome-wide rate (Supplemental Table S2), and shifted permutation Z-scores reveal that the enrichment is specific to the ATAC-Seq peaks as opposed to wider genomic regions (Figure 1). We also considered 6,013 deletions (with 7,365 unique breakpoints) that were present in the DDD consensus dataset (Deciphering Developmental Disorders Study 2015) (Methods) at a frequency of at least 1%, representing variants expected to be relatively common in human populations (Supplemental Table S1). These variants show a dip in frequency and downward trend near active sites (Figure 1a). However, we note that the overlap between common variant breakpoints and ATAC-Seq peaks is still ∼ 2-fold higher than the expected genome-wide rate (p < 10^-4^). We conclude that singleton deletion breakpoints often occur at TFBSs in spermatogonia, suggesting a higher mutational input or less accurate repair at these sites compared to neighbouring regions. The breakpoints of more common variants are observed less frequently at the same binding sites, which may indicate the action of purifying selection in the removal of deleterious mutations at these active regulatory sites.

**Figure 1:**
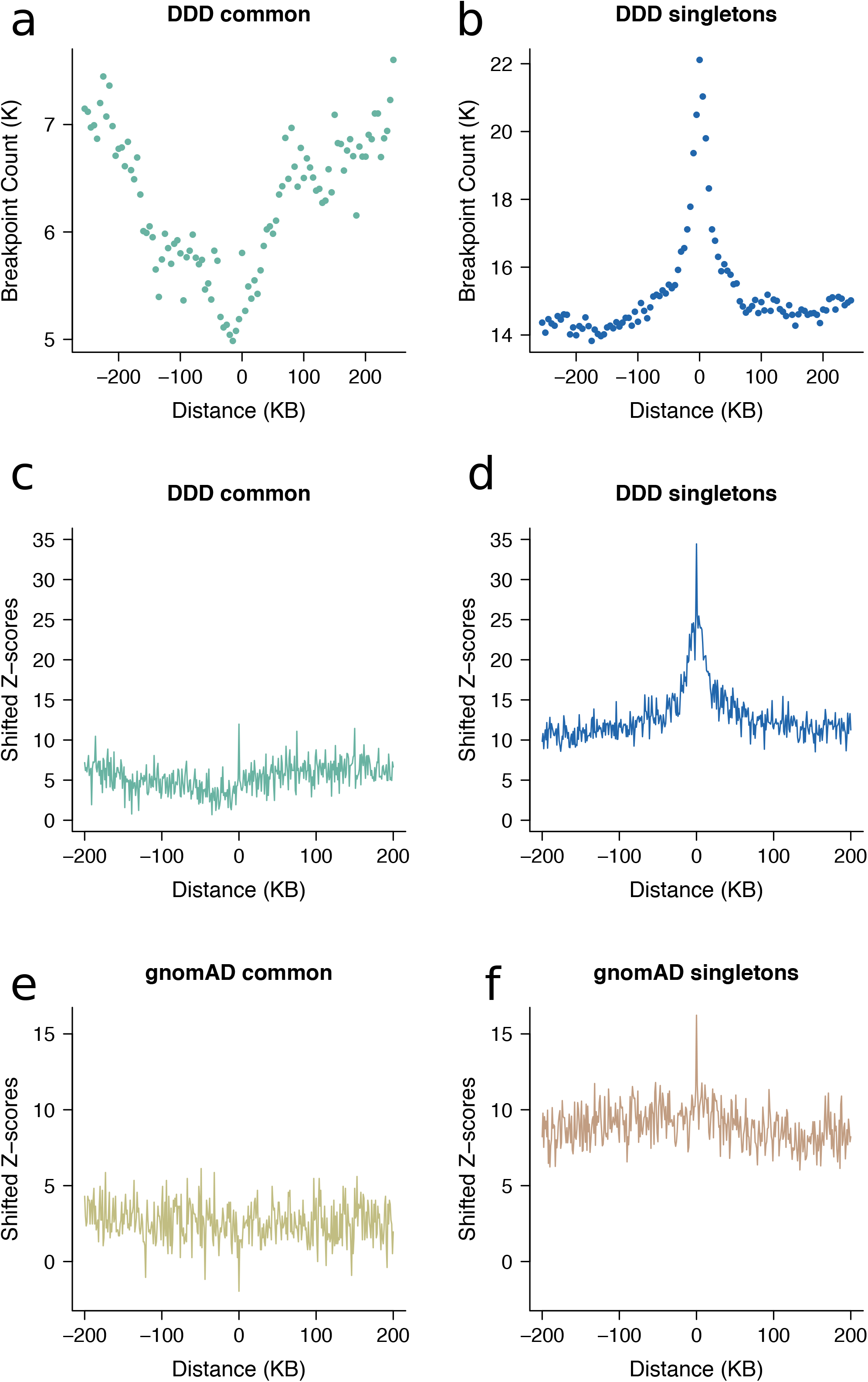
Locally elevated structural variation rates at spermatogonial regulatory sites. SV breakpoint count (a, b) and circular permutation shifted Z-scores (c, d) of deletion breakpoints in the DDD cohort, centred around the midpoints of spermatogonial ATAC-Seq peaks. “Singletons” are breakpoints of deletions with a frequency of ∼ 0.002% across population samples; “common” variants are seen at a frequency of at least 1% in the DDD consensus dataset (see main text); permutation p-values indicate significant enrichment for both types of variants at ATAC-Seq peaks (p < 10^-5^ in each case) (e, f) Circular permutation shifted Z-scores of gnomAD deletion breakpoints, centred around spermatogonial ATAC-Seq peaks. “Singletons” are breakpoints of deletions with a frequency of ∼ 0.002% across population samples; “common” variants are seen at a frequency of at least 5% in the gnomAD V.2 dataset. Permutation p-values indicate significant enrichment for singleton breakpoints (p < 10^-5^), and a significant depletion for common variants (p < 0.01).

Similar trends are also observed for singleton deletion breakpoints from an independent large-scale aggregated dataset of human variants (Figure 1e, 1f) from whole genome sequence (WGS) analysis (Collins et al. 2020) (Supplemental Table S1). We again find a significant enrichment of singleton variant breakpoints at ATAC-Seq peaks, and this enrichment is not seen for common variants (Figure 1).

### Locally elevated mutation at spermatogonial TFBSs

Compared to larger structural variants, such as those (up to megabase sized) deletions examined above, indels have been shown to occur at a higher rate of about 6 new variants per genome and generation (Besenbacher et al. 2016). Short indels (<= 4 bp) are thought to arise due to replication slippage (Levinson and Gutman 1987; Montgomery et al. 2013), whereas longer variants have been considered a hallmark of inaccurate DNA repair after DSBs (Rodgers and McVey 2016). Here, we focus on gnomAD singleton indels <= 20 bp as these variants are expected to be well resolved using short read sequencing. To enable higher spatial resolution of the mutation patterns at ATAC-seq defined accessible chromatin regions, and for the subsequent inference of the associated DNA binding proteins, we identified 706,008 protein binding sites using ATAC-Seq footprinting analysis (Li et al. 2019) (Methods; Supplemental Tables S3 and S4). The rate of singleton 5-20 bp insertions at footprinted spermatogonial protein binding sites approximately doubles from background expectation and is highly concentrated to within 1 kb of the binding site (Figure 2); shifted Z-scores based on genome-wide circular permutations similarly show a highly localized spike of insertions around TFBSs (Figure 2). This pattern starkly contrasts the localised depletion of common variants of the same mutation class at the same binding sites (Figure 2), again implicating a locally elevated mutation rate and purifying selection. In fact, most classes of rare mutation (singleton SVs, smaller and longer indels, SNPs) are significantly enriched at spermatogonial TFBSs (Figure 3), and in the gnomAD dataset, where all singleton classes have been ascertained by WGS, the enrichment is strongest for insertions >= 5 bp. We confirmed the enrichment of singleton short insertions and SV deletion breakpoints at spermatogonial TFBSs, using an independent permutation approach with bedtools shuffle (Quinlan and Hall 2010) (Supplemental Table S5).

**Figure 2:**
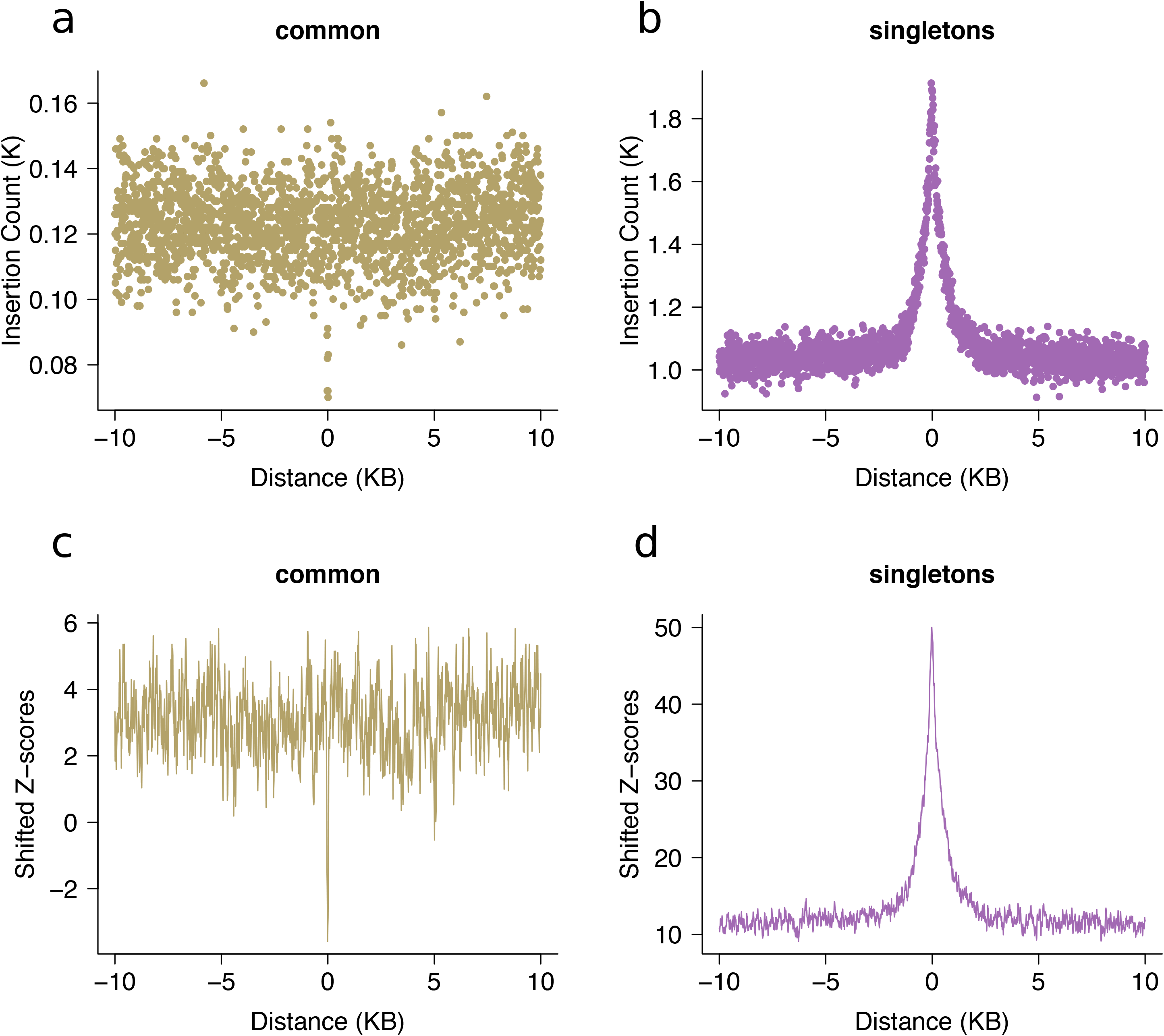
Increased rates of short insertions focussed on spermatogonial binding sites. Insertion count (**a, b**) and Shifted Z-scores (**d, e**) of gnomAD singleton and common insertions (5-20 bp), centred around spermatogonial TFBSs. Singletons are seen only once in the gnomAD V.3 dataset (allele frequency <= 0.001%) and are significantly enriched at binding sites (p < 10-4); common variants have an allele frequency of at least 5% within gnomAD V.3 and are significantly depleted at binding sites (p < 10-4).

**Figure 3:**
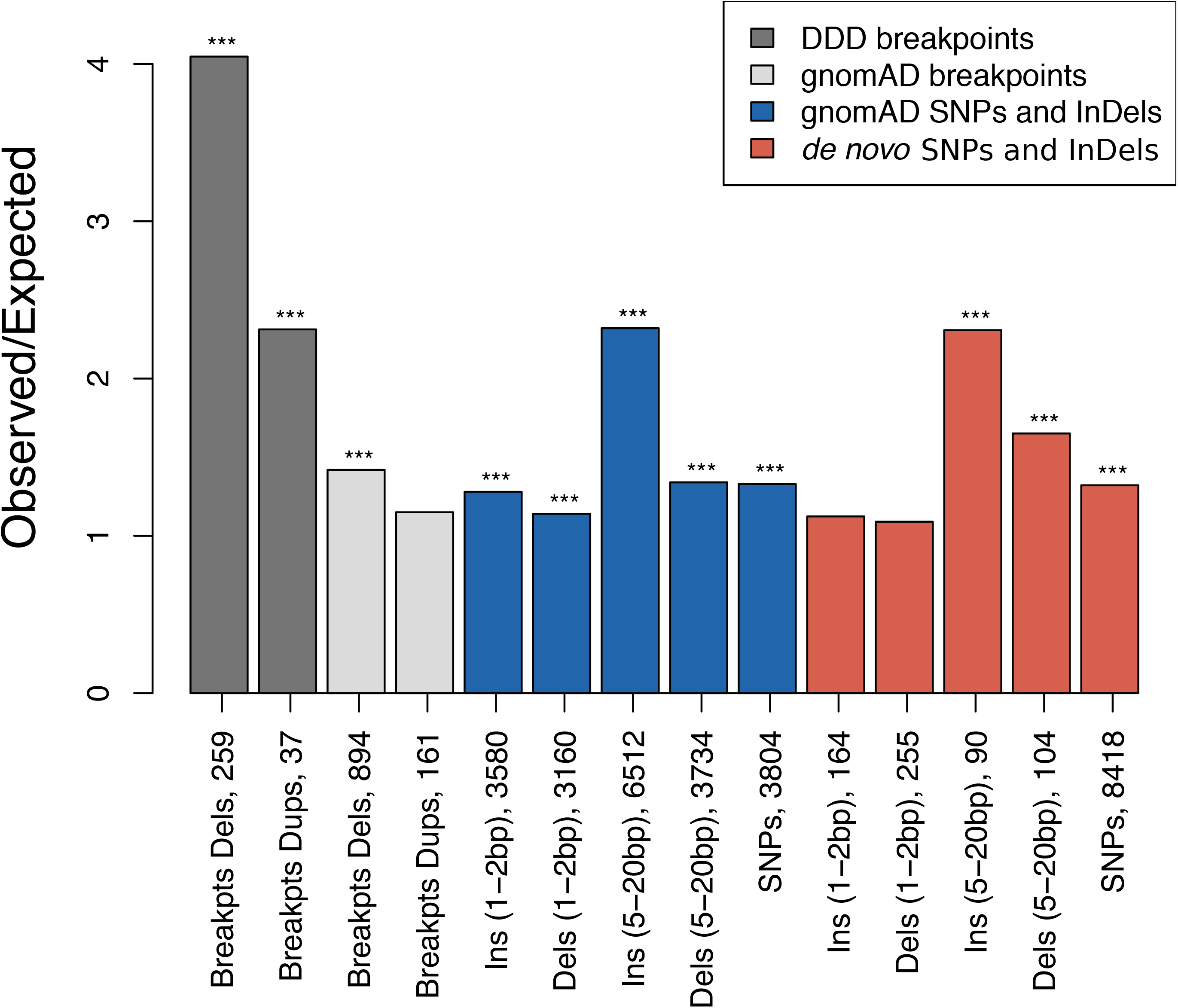
Parallel enrichments of short variants and SV breakpoints at spermatogonial binding sites. Circular permutation results are based on 10,000 permutations; results for singleton variants and de novo mutation are shown. The Y axis shows the ratio of observed over expected variant counts. Mutation categories with significant enrichment are indicated by asterisks (*** indicating p < 0.001). The type of variant tested and the total number of observed variants overlapping spermatogonial TFBSs are indicated below each bar.

In addition to singleton variants from large population samples, we also compiled a set of “gold standard” *de novo* short variants from a range of trio sequencing studies (see Methods). The *de novo* variants show a strikingly similar trend to the gnomAD singleton variants, with a moderate (∼10-60%) increase of mutation rates at TFBSs for all categories of short 1-2bp sequence variants, a but larger increase of ∼130% for insertions of 5-20 bp (Figure 3). These results were confirmed using a set of independent positive and negative control sites (Supplemental Fig. S3). We conclude that regulatory sites that are active in spermatogonia show unusual parallel enrichments for both large SV breakpoints and 5-20 bp insertions, consistent with localised DNA damage or error-prone repair.

### Germline PRDM9 and NRF1 binding generate hotspots for structural variation

To examine any differences in mutational loads associated with different binding factors, we analysed mutational patterns stratified by the binding factors included in the JASPAR database (Sandelin et al. 2004). We accounted for redundancy caused by multiple factors binding to a single motif by considering 167 motif families (Supplemental Table S6). Furthermore, using the reported binding site motif for PRDM9 (Myers et al. 2008), we defined 9,778 putative PRDM9-bound sites corroborated by evidence for H3K4me3 enrichment in testis (Methods). The spermatogonial binding sites of 11% (19/167) of motif families overlapped DDD singleton deletion breakpoints more often than expected (Bonferroni corrected *p* = 0.017) and no motif family was found to be depleted for breakpoints (Supplemental Table S3), suggesting that increased load is a common feature of TFBSs bound by different transcription factors in the germline. Similarly, singleton 5-20bp insertions from the gnomAD database were found to be significantly enriched at 29% (48/167) of families (Bonferroni corrected p = 0.017) and, nominally, 84% (140/167) of families showed enrichment for these insertions (Supplemental Table S4). Again, no TFBS family was found to be depleted for these rare variants. Collectively, these results suggest that TFBSs active in spermatogonia incur locally elevated burdens of short insertions and large structural variants across many different binding motifs.

Certain motif families appear to carry notably higher mutational loads than the general disruption seen across all TFBSs. Based on the insertion fold enrichment (IFE), i.e. the ratio of the observed to expected numbers of insertions (5-20 bp), PRDM9 binding sites are among the most disrupted sites in the genome (IFE = 6.3), and this also holds for PRDM9 sites outside known sites of recombination (IFE = 6.7 for 8,139 PRDM9 sites with a distance of at least 500bp from HSs and ssDNA sites, respectively). PRDM9 sites are similarly associated with higher rates of singleton deletion breakpoints (Figure 4a, 4c), in line with the roles of PRDM9 during recombination, though PRDM9 sites outside known sites of recombination also show this trend (observed overlaps with deletion breakpoints = 9; expected = 1; *p* < 10^-4^). Two other TFBS families, corresponding to NRF1 (Nuclear Respiratory Factor 1; IFE=7.0) and HINFP (IFE=6.6) exceed the disruption seen at PRDM9 sites, and remarkably, NRF1 sites are disrupted at high rates according to DDD breakpoint data (Supplemental Table S3). Shifted Z-scores for the enrichment of short insertions (5- 20 bp) at both NRF1 and PRDM9 binding sites are in the top four, next to SP/KLF transcription factors (motif families 938 and 992), suggesting strong focal enrichments at these sites (Supplemental Tables S4 and S6). NRF1 has been shown to be an important testis-expressed gene with meiosis-specific functions (Wang et al. 2017; Palmer et al. 2019), but NRF1 binding sites have, to our knowledge, not been reported to be foci for genomic instability. We find strikingly similar enrichments of short insertions (5-20 bp) at TFBSs in SSEA4- and KIT-marked spermatogonial samples produced in previous ATAC-seq studies (Guo et al. 2017; Guo et al. 2018). Reprocessing these previous datasets identically to our own reveals that PRDM9, NRF1 and HINFP sites are again among the top 5 disrupted motif families (Supplemental Tables S7 and S8).

**Figure 4:**
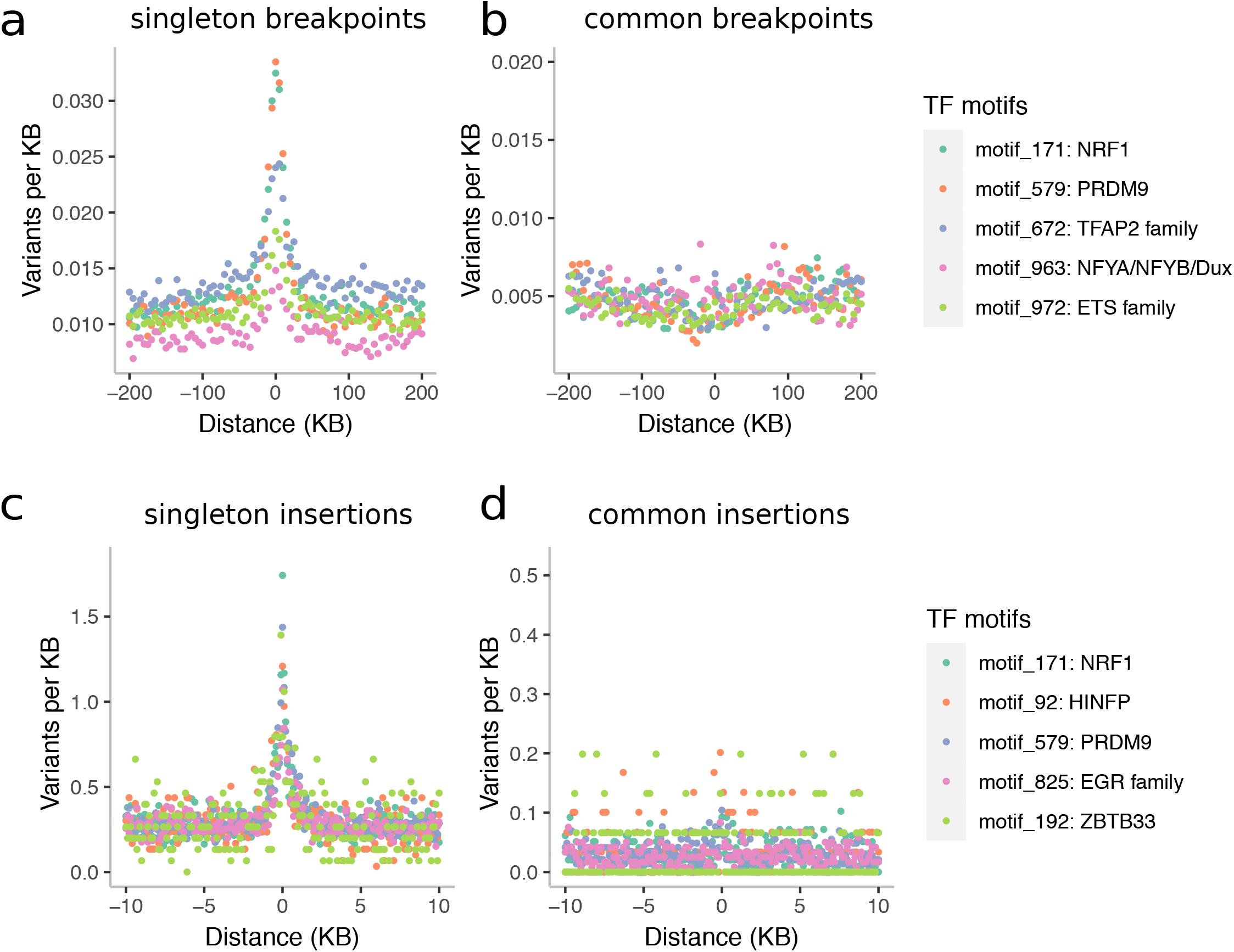
Binding factors associated with the highest rates of mutation at spermatogonial binding sites. Plots are centred on the binding sites of a given motif family inside ATAC-Seq footprints. (**a**) Singleton and (**b**) common deletion breakpoints in the DDD cohort; singletons are breakpoints of deletions with a frequency of ∼ 0.002 across population samples; common variants are seen at a frequency of at least 1% in the DDD consensus dataset. (**c**) Singleton and (**d)** common insertions (5-20 bp) in the gnomAD dataset. Singletons are seen only once in gnomAD V.3 (allele frequency <= 0.001%), and common variants have an allele frequency of at least 5% within gnomAD V.3. Only 10 kb regions around TFBSs with >=95% unique mappability (umap24 scores) were included. The top 5 disrupted motifs are shown, listed in order of enrichment of singleton variants in the circular permutations (all enrichments of singletons are associated with p-values < 10^-4^).

Although both PRDM9 and NRF1 binding sites are GC-rich, their modest motif similarity suggests that the two factors occupy distinct binding motifs (PWMclus: Pearson’s correlation distance *r* = 0.35 for PRDM9 *versus* and NRF1) and should not converge on the same sites. However, in practice, PRDM9 and NRF1 binding sites were often found within the same regulatory regions, such that many (1,199) ATAC- Seq peaks contained both the NRF1 and PRDM9 binding motifs. The disruption of motifs within these co-bound peaks was notably higher, with NRF1-motifs being disrupted by short insertions 10.8-fold the expected rate (observed: 108; expected: 10), and PRDM9-motifs 11.2-fold the expected rate (observed: 146; expected: 13) when co-occurring with the other factor (*p* < 10^-4^ in each case). Similarly, 1,311 ATAC-Seq peaks contained a motif for both CTCF and PRDM9, and CTCF motifs in these peaks were more highly disrupted by short insertions (ratio = 6.3; observed: 69; expected: 11) compared to all CTCF motifs (Supplemental Table S4), as was PRDM9 (ratio = 8.2; observed: 115; expected: 14) (*p* < 10^-4^ in each case). Importantly, the excess of insertions observed at particular motif sites is not a trivial consequence of statistical power (i.e. the number of TFBSs in the genome); for example, the number of binding sites identified for PRDM9 and NRF1 is fewer than many other factors (< 10,000 sites each; Supplemental Tables S3 and S4). In general, mutational loads appear to be dependent on the level of chromatin accessibility (MACS2 peak scores) and the number of factors predicted to bind at ATAC-Seq defined regulatory regions, such that regions in the upper quartile of accessibility that are also occupied by more than 4 factors incur the highest indel loads (Supplemental Fig. S4). The significant positive correlation between the rates of binding site disruption via singleton insertions and deletion breakpoints across all motif families (Supplemental Fig. S5; Spearman’s R = 0.52, p < 10^-5^) suggests that the two types of damage may be mechanistically linked. In support of this idea, singleton short insertions (5-20 bp) and singleton SV deletion breakpoints overlap at the exact nucleotide position more often than expected (genome-wide Z-score = 26.31; *p* < 10^-4^; see also Supplemental Fig. S6). This overlap is unlikely to be due to erroneous variant calling in the singleton dataset since we observe similar patterns for common variants of the same variant categories (genome-wide Z-score = 62.9, *p* < 10^- 4^).

### Short insertions generate clustered binding sites within regulatory regions

Intriguingly, the 5-20 bp insertions observed at TFBSs frequently occur within only a few nucleotides of the binding motifs, whereas other classes of short variants do not show such a precisely localized increase (Figure 5 and Supplemental Fig. S7). Despite a moderate genome-wide enrichment (Figure 3), the 1-2 bp insertions characteristic of polymerase slippage, do not peak in the immediate neighbourhood of TFBSs (Figure 5 and Supplemental Fig. S7). We examined the consequences of elevated 5-20 bp insertion rates at TFBSs using an exhaustive motif search algorithm (Bailey et al. 2009), which finds overrepresented sequence motifs among a set of input sequences. We found that the inserted sequences at a mutated TFBS often contain additional copies of the sequence motif corresponding to the original TFBS (Figure 6 and Supplemental Fig. S8), suggesting that many insertions at TFBSs are tandem duplication events, including events at CTCF, NRF1 and PRDM9 sites. The presence of these motif-containing singleton insertions appears to reveal a novel mutational mechanism expected to increase the number of binding sites for a binding factor and to lead to the expansion of TFBS clusters. CTCF-binding sites are known to occur in clusters (Kentepozidou et al. 2020) and are often affected by singleton insertions in our dataset (ranked 12^th^ out of 167 motif families, based on the number of insertions per TFBS; Supplemental Table S4). We find that spermatognial active sites exhibit greater homotypic clustering of TFBS than ATAC-seq defined binding sites from somatic tissues (Figure 6). Combined with a positive correlation between homotypic motif clustering and insertion rate, this suggests that spermatogonia binding sites are progressively accruing motif clusters.

**Figure 5:**
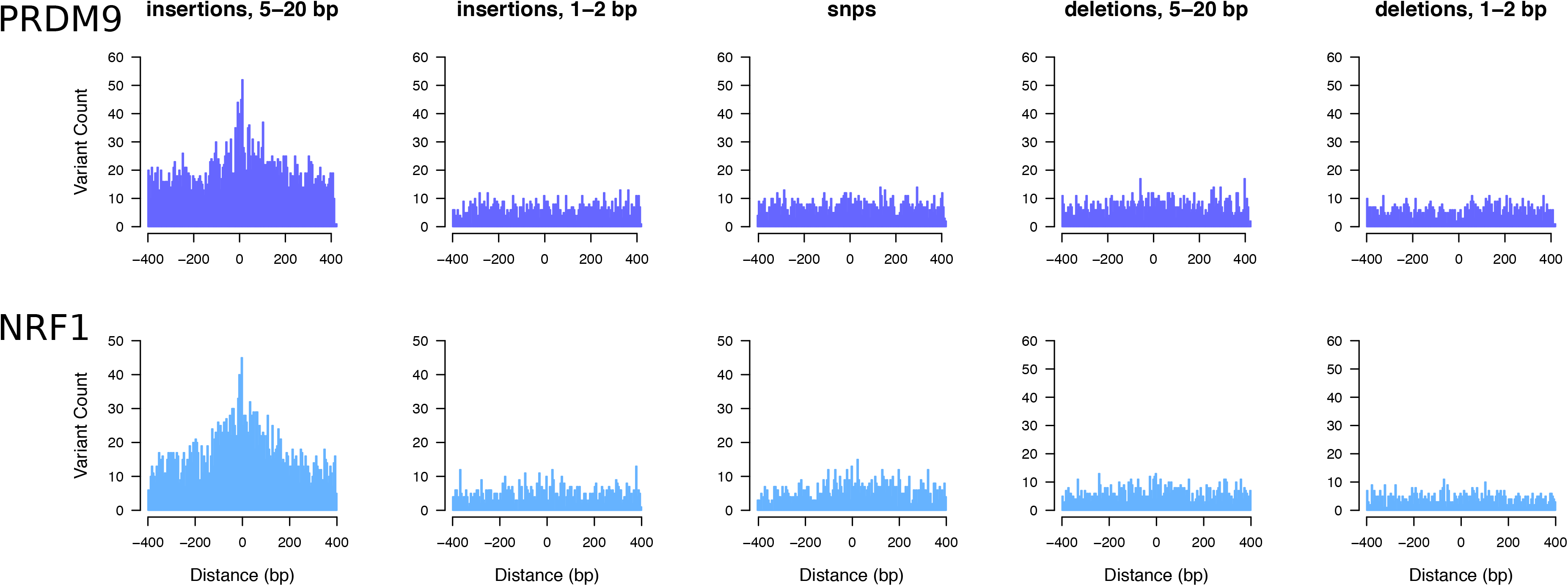
Elevated singleton insertion rates at PRDM9 and NRF1 binding sites contrast with other short variant classes. All gnomAD variants have been down- sampled to a total of 650,000 variants per analysis, making the Y axes directly comparable; individual bins are 5 bp in size. Only regions around TFBSs with >=95% unique mappability (umap24 scores) were included.

**Figure 6:**
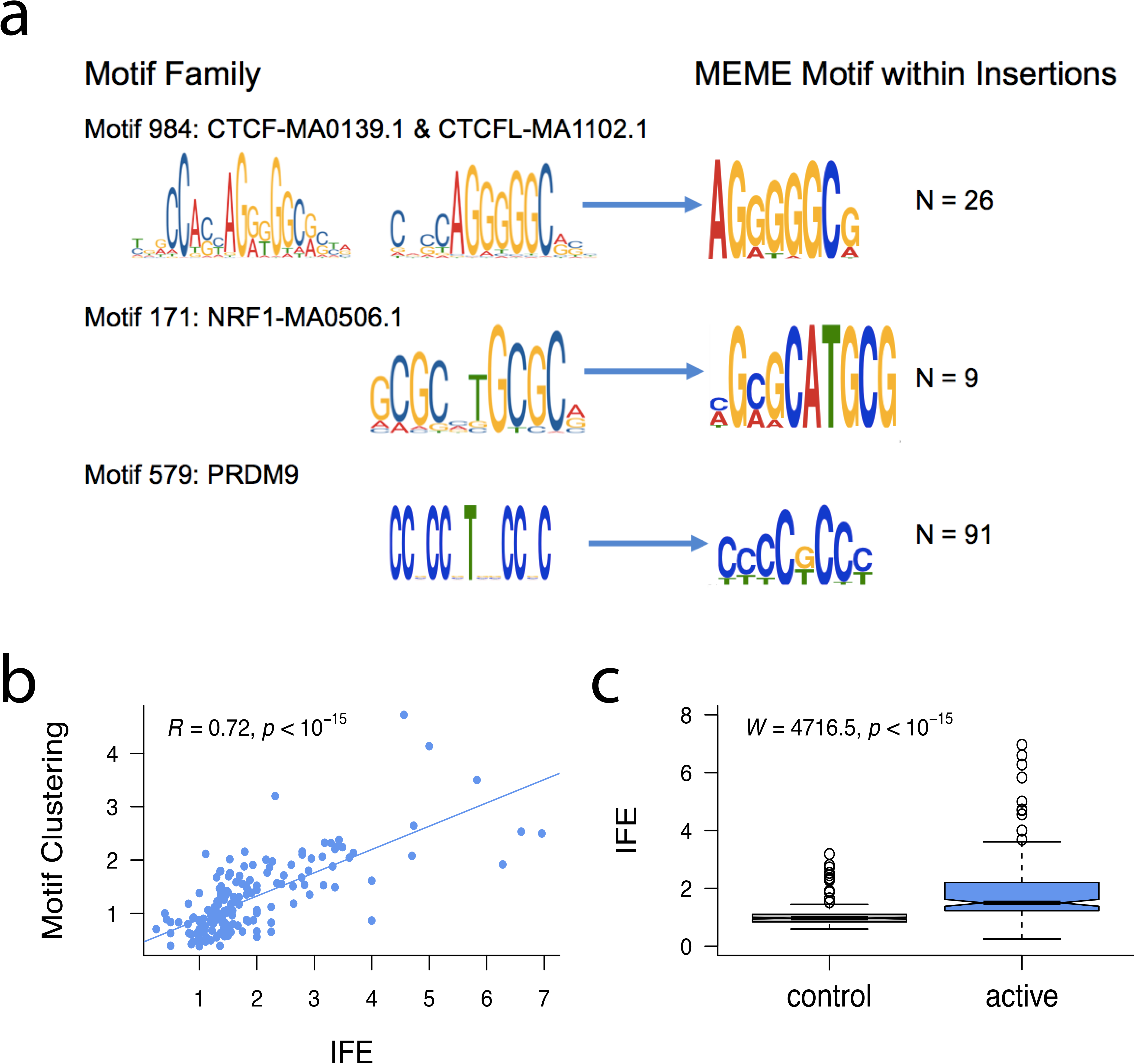
Insertions at spermatogonial TFBSs generate motif clusters in the genome. a) Jaspar database sequence motifs identified in the footprints of spermatogonial ATAC-Seq peaks (left) and the motifs identified in the singleton insertions (5-20 bp) (right). The number of insertion sites (N) that were chosen by MEME to construct the motif are shown on the right. **b)** For each motif family, we plot the insertion fold enrichment (IFE) on the X axis and the degree of spermatogonial motif clustering on the Y axis; the least square regression line is indicated in blue. Motif clustering is measured as the distance to the nearest motif at spermatogonial active sites, relative to the distance for motifs at ENCODE active sites. c) The insertion fold enrichment (IFE) is contrasted between FIMO control motif sites (left) and spermatogonial active motif sites (right); the Wilcoxon Test was performed to compare the IFE at the two classes of sites.

These unusual patterns of clustered TFBSs at indel breakpoints appear to be specific to spermatogonial ATAC-seq peaks, and do not reflect genome-wide trends. Applying the MEME-Chip algorithm on 50bp regions flanking singleton insertion and deletion breakpoints, we were able to re-discover the sequence motifs of commonly disrupted binding sites, including the motifs of PRDM9 and NRF1 (Supplemental Table S9). In contrast, genome-wide, the motifs discovered flanking these variants were more likely to be simple repeats and other low complexity sequences that did not match known TFBS motifs, suggesting that processes other than transcription factor binding drive DNA breakage outside of active regulatory sites.

### Genomic instability at spermatogonial TFBSs impacts enhancers active in neural development

Since many regulatory regions of the genome are active across a variety of cell types (Andersson et al. 2014), mutation at TFBSs in spermatogonia might disrupt gene regulation in other tissues. The developing brain is of particular interest, given reports of increased SV burdens in neurodevelopmental disorders (Girirajan et al. 2011; Leppa et al. 2016; Collins et al. 2017). We classified developmentally active human brain enhancers (distal regulatory elements) supported by neocortical ATAC-Seq data (de la Torre-Ubieta et al. 2018) according to whether they were either active (10,888 brain enhancers) or inactive in the male germline (26,162 brain enhancers). We then calculated the odds ratio of a singleton mutation affecting a brain enhancer which is also *active* in spermatogonia, relative to a brain enhancer which is *inactive* in spermatogonia. For DDD singleton deletion breakpoints, the odds ratio was 6.82 (95% CI = [5.34,8.71]), and for a singleton gnomAD insertion (5-20 bp), it was 4.69 (95% CI = [4.46,4.93]). This suggests that activity in spermatogonia greatly predisposes a brain enhancer to DNA damage, and this damage manifests in enhancers that share activity with the male germline (Figure 7). Brain enhancers that are shared with spermatogonia are, on average, more accessible in the developing brain than those that are inactive in the germline (the median “mean of normalized counts” for the two types of brain enhancers were 104.8 and 54.1, respectively; Wilcoxon test W = 197340000, p-value < 2.2e-16), suggesting a link between enhancer activity, the sharing of enhancers across tissues and propensity to mutation. The subset of brain enhancers which overlapped spermatogonial active sites were not enriched for specific motifs, and the number of motif sites for each motif family were highly correlated between brain and spermatogonia (Spearman’s rho = 0.95, *p* < 10^- 15^). That is, the propensity to mutation does not appear to be driven by an enrichment of specific motif families in brain enhancers.

**Figure 7:**
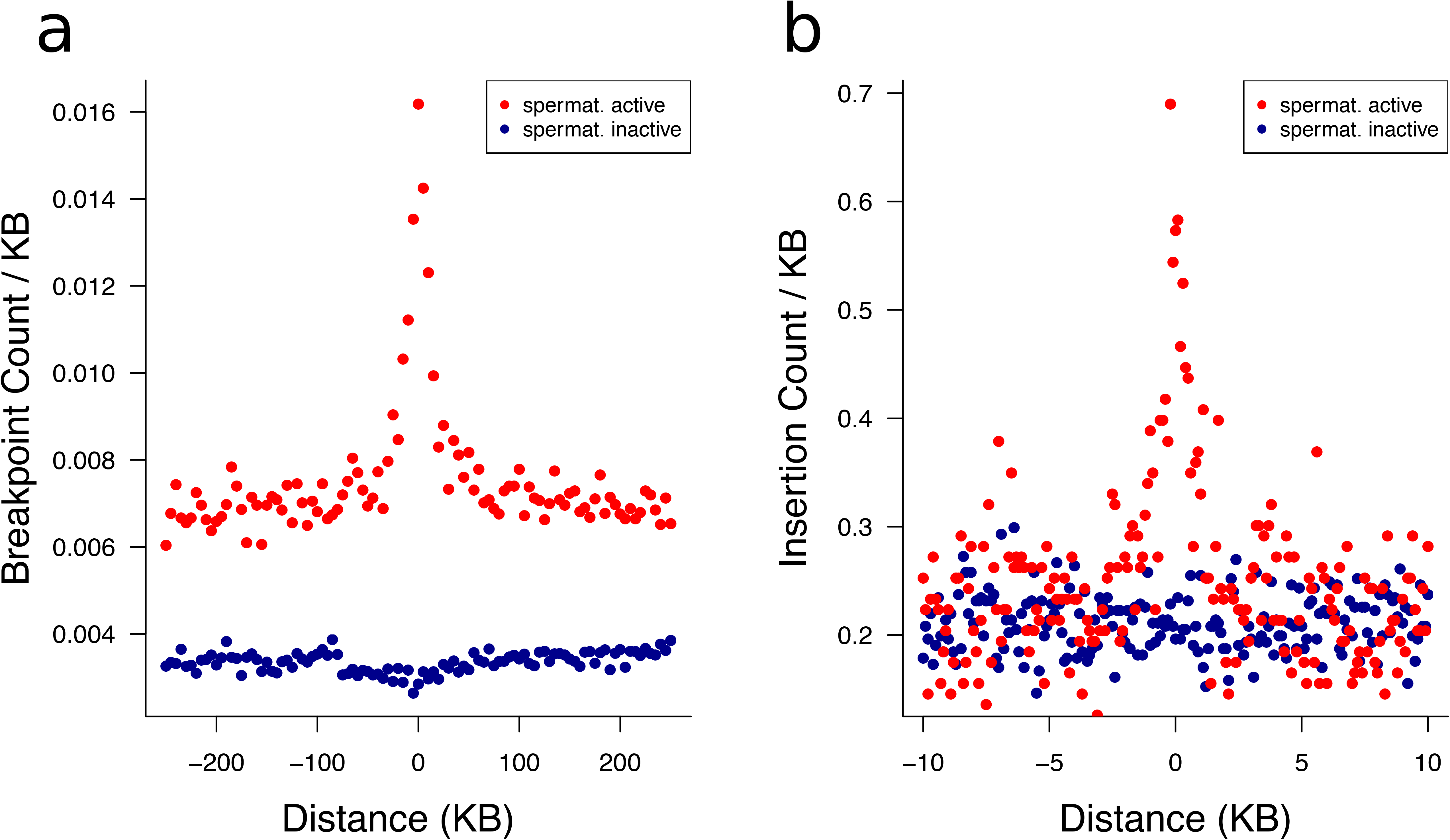
Neural enhancers with activity in spermatogonia suffer elevated mutation rates. **a)** Singleton DDD deletion breakpoint and **b)** singleton gnomAD insertion (5- 20 bp) count around brain active enhancers. Enhancers were classified as being also active in spermatogonia (red) or inactive in spermatogonia (blue). Plotted is the average number of variants per brain enhancer - in 5 kb windows or 100 bp windows, respectively. In **b**, only 10 kb regions around enhancers with >=95% unique mappability (umap24 scores) were included (3,409 brain enhancers that are inactive in spermatogonia and 1,029 that are active).

### Spermatogonia accessible TFBS motifs incur increased rates of disruption

We cannot exclude a small contribution of the TFBS sequence itself on the predisposition to mutation (Kondrashov and Rogozin 2004), but our data suggest that TF binding is a major driver of insertion and deletion mutation in the human germline. This is supported by the fact that we see an increase of disruption of brain enhancers if they are active in spermatogonia (Figure 7) and, more generally, an increase in the mutational load for sites that are active across other somatic tissue if binding also occurs in the germline (Supplemental Table S10). In addition, control motif sites (representing the same TFBS but located outside of ATAC-Seq peaks) are subject to lower rates of mutation compared to motifs within spermatogonial ATAC- Seq peaks (Figure 6c). Motifs within peaks carry, on average, 73% more mutations than their control counterparts, and for the most highly disrupted motifs, the discrepancy between active and control motifs is even larger. For example, PRDM9 motifs are 3.4-fold, HINFP 2.9-fold and NRF1 motifs 2.6-fold more disrupted if they are active in spermatogonia, relative to spermatogonia inactive motifs. We note that this increase in disruption is likely to be a conservative estimate since some control sites may be bound at time points in the germline that our ATAC-Seq data cannot ascertain.

Since the X chromosome spends only one third of its time in males - the sex with the higher number of germ cell divisions - a depletion of mutations on the X chromosome is expected for a male-biased mutational process. We find the X chromosome to be strongly depleted for short singleton gnomAD insertions (5-20 bp), with a ratio of X to autosome variants per uniquely mappable site of 0.78 (Supplemental Table S11). However, we note that, despite the overall reduced rate of insertions on the X, ATAC- Seq peaks on the X are still subject to increased rates of insertions compared to genome-wide expectations, suggesting that the inferred effects of protein-binding on mutation are larger than the reduction in mutation due to X-linkage (38 observed insertions in X-linked ATAC-Seq peaks, whereas 11 were expected; p < 10^-4^).

To test which candidate genomic feature most reliably predicts DNA damage, we used random forest regression to model the rate of singleton variants within 5 kb genomic windows, based on their overlap with spermatogonial TFBSs, ssDNA sites, LD-based hotspots, average GC content, mappability, gene density, replication time as well as various repeat families (LTRs, SINEs, LINEs and simple repeats). In models of genome-wide short insertion rates or deletion breakpoint rates, measures of replication timing and GC content were important predictors of mutation load as expected (Supplemental Fig. S9). Mappability was an important factor for predicting mutation rates genome-wide, perhaps reflecting the association between segmentally duplicated (low mappability) regions and rapid structural evolution, or perhaps suggesting that a fraction of variants may be erroneously called in the gnomAD dataset. (Only regions with high mappability were included in our more detailed analyses of TFBSs (Figures 3-7 and Supplemental Fig. S7)). However, spermatogonial ATAC-Seq derived TFBSs contributed additional predictive power to the models, even at the scale of the entire genome. The same TFBSs appear to be somewhat more important features in models that specifically predict damage at active brain enhancers (Supplemental Fig. S9). Genome-wide, deletion breakpoints and 5-20 bp insertions were enriched in early replicating DNA (Spearman’s rank correlation with replication timing: rho = 0.08, p < 10 ^-15^ and rho = 0.07, p < 10^-15^, respectively). In contrast, the presence of repeat elements had almost no impact in predicting either short insertion or deletion breakpoint rates (Supplemental Fig. S9). We conclude that germline active regulatory sites, through their occupancy by DNA binding factors, make a substantial contribution to genome-wide *de novo* structural variant rates, independent of other genomic features.

## Discussion

We have demonstrated enrichments of rare and *de novo* SV breakpoints at spermatogonial regulatory sites defined by ATAC-Seq, suggesting that these sites suffer high rates of DSBs in the male germline. The same sites show unusual parallel enrichments for short variants, and particularly 5-20bp insertions. No TFBS family examined was found to be depleted for these rare variants, suggesting that many different TFBSs active in spermatogonia are prone to higher mutational loads. These loads appear to be positively correlated with the levels of chromatin accessibility/nucleosome disruption (ATAC-Seq peak binding strength) and the number of factors predicted to bind within the region. Sites bound by PRDM9, NRF1 and HINFP incur the highest levels of disruption, but 11% of 167 TF families examined showed evidence for significantly elevated mutation rates. These results have implications for the evolution of binding site patterns within regulatory regions, and for disrupted regulation in somatic tissues.

Homotypic clusters of TFBSs are a pervasive feature of both invertebrate and vertebrate genomes, and have long been known to be a common feature of human promoter and enhancer regions (Gotea et al. 2010). Various adaptive hypotheses have been proposed for the presence of such clusters such that they provide functional redundancy within a regulatory region, enable the diffusion of TF binding across a region, and allow cooperative DNA binding of TF molecules (Gotea et al. 2010). More recently it has been suggested that homotypic TFBS clusters may also contribute to phase separation and the compartmentalisation of the nucleus (Kribelbauer et al. 2019). Similarly, the clustered patterns of CTCF sites in the genome have been ascribed critical roles in chromatin architecture and regulation, particularly at regulatory domain boundaries. However, these boundary regions have been shown to exhibit genome instability (Kaiser and Semple 2018) and recurrently acquire new CTCF binding sites in dynamically evolving clusters (Kentepozidou et al. 2020). The data presented here suggest that binding site clusters may arise solely as a selectively neutral consequence of the unusual mutational loads at germline TFBSs, with clusters maintained by recurrent DNA damage and misrepair.

We observe significant enrichments of both large SV breakpoints and small insertions together at spermatogonial TFBSs. This parallel enrichment of both types of mutation may originate from DNA breakage, followed by misrepair, conceivably via a pathway such as non-allelic homologous recombination (NAHR). It is known that NAHR can create large insertions and deletions (Kim et al. 2016), and PRDM9 activity is implicated in certain developmental disorders arising via NAHR (McVean 2007; Myers et al. 2008; Berg et al. 2010). For example, the locations of PRDM9 binding hotspots coincide with recurrent SV breakpoints causing Charcot-Marie-Tooth disease, and Hunter and Potocki-Lupski/Smith-Magenis syndromes (Pratto et al. 2014). It is possible that the sequence similarity at TFBSs scattered across the genome may make them particularly prone to NAHR. However, we note that the sequence similarity between the low copy repeat units, known to be involved in NAHR, is usually of the size of several kb (Gu et al. 2008), rather than sequences on the scale of TFBSs. The NHEJ pathway can also lead to short insertions after DNA breakage, usually in G0 and G1 phases of the cell cycle. Indeed, NHEJ is the most common repair pathway of DSBs in mammals and it is typically error prone (van Gent et al. 2001; Lieber et al. 2003). During NHEJ, double-strand break ends are resected to form single-stranded overhangs, but when pairing occurs between the tips of the overhangs, sequences near the breakpoints will often be duplicated (Rodgers and McVey 2016). Interestingly and consistent with our results based on ultra-rare sequence variants, two previous studies using human–chimpanzee–macaque multiple alignments have shown that high numbers of short insertions have occurred in the human lineage (Kvikstad et al. 2007; Messer and Arndt 2007), and both conclude that these insertions preferentially take place in the male germline, evidenced by decreased mutation rates on the X chromosome, with similar observations in rodents (Makova et al. 2004).

The data presented here suggest that different DNA binding proteins differ widely in their impact on mutation rates. The two proteins with the largest impacts, NRF1 and PRDM9, are both highly expressed in testis, revealing a possible link between the expression level of a gene encoding a DNA binding protein and the propensity for breakage or inefficient repair at the sites the protein binds. Incidentally, NRF1 has a pLI score of 0.999, indicating that it is extremely loss-of function intolerant and crucial for the organism’s functioning (Karczewski et al. 2020). A previous study (Montgomery et al. 2013), using 1K genomes polymorphism data, failed to find an increase in indels at PRDM9 motifs genome-wide. This highlights the importance of using ATAC-Seq data to confine the search for motifs to germline active sites only, combined with singleton variants from large-scale sequencing studies as a more powerful strategy to explore fine scale mutational patterns.

Although studies of coding sequences, such as the DDD (Deciphering Developmental Disorders Study 2015), have revealed many of the genes disrupted in developmental disorders, more than half of cases lack a putatively causal variant (Mcrae et al. 2017), stimulating interest in the noncoding remainder of the genome, and particularly regulatory regions active in development. Limited sequencing data, covering a fraction of human regulatory regions, suggests that *de novo* mutations are enriched in these regions and are therefore likely to contribute to neurodevelopmental disorders at some level (Short et al. 2018; Gerrard et al. 2020). However, there appear to be very few, if any, individual regulatory elements recurrently mutated across multiple cases to cause neurodevelopmental disorders with a dominant mechanism (Short et al. 2018). The data presented here suggests a potential solution to this paradox, where combinations of mutations at multiple regulatory regions may underlie a disease phenotype. The frequency of such combinations is expected to be many times higher if they involve regulatory regions bound by factors such as NRF1. In such cases, an entire class of sites, rather than an individual site, is subject to recurrent mutation.

## Methods

### Identification of spermatogonial binding sites

Samples of testicular tissue were obtained from three patients undergoing orchiectomy with total processing completed within ∼5-7 hours of explant. Tissue was obtained after informed consent through the Lothian NRS BioResource, and the study was approved by NHS Lothian (Lothian R&D Project Number 2015/0370TB). Tissue samples were disaggregated into cells, and cells were labelled with phycoerythrin (PE)-conjugated antibody against the cell surface marker FGFR3 (FAB766P, clone 136334, R&D systems). Spermatogonial cells were isolated using a FACSAria II cell sorter (BD bio- sciences) based on PE fluorescence and cell shape, according to Forward/Side Scatter. Isolated cells were subjected to ATAC-seq using the protocol and reagents described in (Buenrostro et al. 2013), followed by paired- end sequencing on Illumina HiSeq4000 (75 bp read length). We combined reads from separate sequencing runs into three biological replicates, based on origin and morphological appearance of the FACS sorted cells. Replicate 1: combined sequencing runs H.5.1 and H.5.4; a non-cancer patient; large cells, high side scatter; 58,000 and 42,000 cells, respectively. Replicate 2: combined sequencing runs H.5.2 and H.5.5; the same non-cancer patient as Replicate 1; large cells; 36,000 and 23,000 cells, respectively. Replicate 3: combined sequencing runs H.7.3 and H.10.2; normal tissue from cancer patients; large cells; 69,000 and 24,000 cells, respectively. ATAC- seq raw reads were trimmed to remove any retained adaptor sequences using cutadapt (Martin 2011) with parameters *-n3 --format=fastq --overlap=3 -g GAGATGTGTATAAGAGACAG -g CAGATGTGTATAAGAGACAG –a CTGTCTCTTATACACATCTG -a CTGTCTCTTATACACATCTC.* Reads were aligned to the GRCh38/hg38 genome assembly with Bowtie2 (Langmead and Salzberg 2012) in paired-end mode, limiting the insert size to 4 kb. Any reads with quality score < 30 were discarded. Paired end reads were converted to fragments using bedtools bamtobed -bedpe, followed by extraction of the most 5’ and 3’ coordinates of each pair. PCR duplicates were removed by retaining only one instance of a fragment with identical coordinates within a sample. Fragments overlapping with the regions previously blacklisted as mitochondrial homologs (Buenrostro et al. 2013) were discarded. Peaks were identified from short fragments of <= 100 bp (Supplemental Fig. S1), thought to arise due to transposition events around transcription factor binding sites – and distinct from fragments spanning the larger nucleosomes (characterized by a ∼200 bp periodicity) (Buenrostro et al. 2013). Peaks were called from short fragments using macs2 callpeak (Zhang et al. 2008) with the following parameters: -B -q 0.01 -f BAMPE --nomodel --nolambda --keep-dup auto --call- summits. The clustering of ATAC-Seq peaks near transcription start sites and promoters was assessed using the ChIPseeker R package (Yu et al. 2015).

For the downstream mutation analyses, ATAC-Seq peaks from Replicates 1 and 2 (the non-cancer patient) were merged, creating a single peak set. This dataset also formed the basis for the footprinting analysis, which used, as input, the combined short sequencing fragments of Replicates 1 and 2, running “rgt-hint footprinting” with --atac-seq and --bias-correction, followed by “rgt-motifanalysis matching” with the option --remove-strand-duplicates (Li et al. 2019). Input motifs were the 579 position weight matrixes (PWMs) of the japspar vertebrate database (Sandelin et al. 2004) as well as the 13-mer PRDM9 motif “CCNCCNTNNCCNC” (Myers et al. 2010) which was also provided as a PWM. The tissue donor for Replicates 1 and 2 was a carrier of the most common (European) alleles of PRDM9, which was confirmed by investigating his allelic state at the SNP (rs6889665) identified by Hinch et al. (2011); this SNP was covered by our ATAC-Seq by 10 reads, all of which were “T”. Accordingly, we assume that the donor is a carrier of the A and/or B allele of PRDM9 (both of which bind the same DNA motif), and the search for the 13-mer PRDM9 motif in this patient’s ATAC-Seq data can be used as a proxy for PRDM9 binding in European populations. In addition, Replicate 3 was processed in the same way as the combined Replicates 1 and 2 and served as a positive control to assess the genome- wide enrichment of mutations at spermatogonial accessible sites (Supplemental Fig. S3).

Jaspar input motifs are often highly similar, resulting in multiple binding proteins being identified by the rgt-hint pipeline to bind at the same ATAC-Seq footprint; this is biologically implausible (since only one protein is likely to occupy a given site), and we clustered motifs by similarity, using the default parameters of the PWMclus CCAT package (Jiang and Singh 2014). This resulted in a set of 167 motif families of similar binding motifs (Supplemental Table S6). Using bedtools (Quinlan and Hall 2010), we merged overlapping binding sites that belonged to motifs of the same family (thus calling them only once), and we also merged palindromic binding sites called on both strands. Since PRDM9 is known to leave a characteristic histone methylation mark on bound DNA (Grey et al. 2011; Powers et al. 2016), we intersected the PRDM9 motif sites with testis-derived H3K4me3 marks (called in an PRDM9 A/B heterozygous individuals) from Pratto et al. (2014). This resulted in a stringent set of PRDM9 sites, which were both located in ATAC-Seq footprints and also carried the H3K4me3 mark in human testis. ATAC-Seq-defined PRDM9 sites showed moderate overlap with DMC1-bound ssDNA sites (Pratto et al. 2014) as well as recombination HSs (Myers et al. 2005), which may reflect the fact that most cells in our experiments are likely to be pre-meiotic: only 10 and 11% of PRDM9 sites were within 500 bp of a ssDNA peak and a recombination HS, respectively, whereas 44% of DMC1-bound sites overlap with LD-defined HSs. However, we find that stronger ssDNA peaks are more likely to be near a PRDM9-binding site (Supplemental Fig. S10).

### Comparisons between ATAC-Seq datasets

Using the same procedure as described above, we processed raw ATAC-Seq reads from previously published datasets in order to call MACS2 peaks from short sequencing fragments. Datasets included ATAC-Seq reads from the germinal zone and cortical plate of the developing brain (SRR6208926, SRR6208927, SRR6208938, SRR6208943) (de la Torre-Ubieta et al. 2018), ATAC-Seq experiments of KIT+ spermatogonia (sra accessions SRR7905001 and SRR7905002) (Guo et al. 2018), SSEA4+ spermatogonia (SRR5099531, SRR5099532, SRR5099533, SRR5099534) (Guo et al. 2017) and ESC cells (SRR5099535 and SRR5099536) (Guo et al. 2017). Adapter sequences within raw sequencing data were identified using bbmerge.sh of bbmap (https://sourceforge.net/projects/bbmap/) and removed using cutadapt (Martin 2011), as above. Encode ATAC-Seq datasets (Encode Project Consortium 2012; Davis et al. 2018) (Liver: ENCFF628MCV, Ovary: ENCFF780JBA, Spleen: ENCFF294ZCT, Testis: ENCFF048IOT, Transverse Colon: ENCFF377DAO) were downloaded as bam files, converted to BEDPE format, and short fragments were identified for peak calling.

BedGraph files (from the MACS2 output), describing the fragment pileup, were converted to bigwig format using bedGraphToBigWig and uploaded to the Galaxy server at https://usegalaxy.eu/ (Afgan et al. 2018). DeepTools2’s multiBigwigSummary (with default parameters) and plotCorrelation (with parameters –skipZeros –removeOutliers) (Ramirez et al. 2016) were used to create a heatmap of ATAC-Seq signals in the different tissues and datasets. The KIT+ and SSEA4+ spermatogonial ATAC-Seq datasets of Guo et al. (2017) and Guo et al. (2018) were further used to perform footprinting and motif matching analyses (Li et al. 2019) as described above for the FGFR3-positive cells. Peaks and motif sites that are accessible in the developing brain but not in spermatogonia were identified using bedtools intersect (Quinlan and Hall 2010).

### Structural Variant Breakpoint data

Large SVs, identified by high-density arrayCGH, or a combination of arrayCGH + exome sequencing, were extracted from a cohort of 9,625 DDD patients, using variant calling procedures as described in (Deciphering Developmental Disorders Study 2015). We filtered the DDD variants to only keep variants which fulfilled the following criteria: a CNsolidate wscore >= 0.468, a callp < 0.01 and a mean log2 ratio of < -0.41 for deletions and 0.36 for duplications; CIFER “false positives” were removed. Singleton variants were identified as being annotated as “novel” by the DDD release, only seen once among the DDD patients, and not seen in the dgv (MacDonald et al. 2014) and gnomAD V.2 (Collins et al. 2020) structural variant datasets (80% reciprocal overlap criterion). Since there are 9,625 patients in the DDD dataset, the gnomAD V.2 dataset contains SVs from 10,738 genomes and the dgv contains SVs from 29,084 individuals, this puts an upper limit of the frequency of carriers of a singleton variant at ∼ 0.002%. Breakpoints were identified as the 5’ and 3’ coordinates of SVs, resulting in 13,406 singleton deletion and 3,406 duplication breakpoints; the resolution of the breakpoints was such that the median and mean confidence intervals were 300 bp and 12 kb, respectively. Further, we identified 11,962 “common” deletion variants in the DDD dataset, which had a minimum variant frequency of 1% in the consensus CNV dataset as described by the DDD study (2015), i.e. pooled CNV datasets of Conrad et al. (2010), the Genomes Project Consortium (2010), the Wellcome Trust Case Control (2010) and the DDD normal controls. We used the 80% reciprocal overlap criterion and grouped common variants using the bedmap options --echo-map --fraction-both 0.8, followed by bedops --merge (Neph et al. 2012). The breakpoints of common variants are thus the outermost coordinates of all SVs that are collapsed into a given variant. The overlap of such “common” breakpoints with ATAC-Seq peaks was assessed independently of SV allele frequencies, i.e. a group of common SVs contributed two breakpoints to the analysis.

We also identified a set of singleton CNVs called with the Manta algorithm (Chen et al. 2016) from the gnomAD V.2 database (Collins et al. 2020) (80% reciprocal overlap criterion with gnomAD V.2, dgv and DDD variants), resulting in a set of 73,063 deletion and 15,419 duplication breakpoints seen in ∼ 0.002% of individuals but called with a different approach compared to the DDD. Common deletions and duplications (*p* >= 0.05) were also extracted from the gnomAD V.2 dataset; these variants had also been called with the Manta algorithm and included 5,954 deletion and 1,586 duplication breakpoint sites.

### Indels and SNP data

The recently released gnomAD V.3 variants (indels and SNPs) were downloaded from https://gnomad.broadinstitute.org/. Only variants that passed all filters were kept (filtering using VCFtools --remove-filtered-all). Multiallelic variants were split using bcftools norm, and bcftools norm --IndelGap 2 was applied to indels, to allow only variants to pass that were separated by at least 2 bp. Singleton variants were defined as having an allele count of one, and the allele number was >=100,000, i.e. the allele frequency of singletons was *p* <= 0.001%.

We subdivided gnomAD indels into singleton insertions and deletions of different sizes: 1-2 bp (most commonly arising due to replication slippage) and those 5-20 bp (arising due to other mechanisms of DNA instability and within the size range reliably detected by short-read sequencing). To speed up simulations and allow for easy comparison between categories of variants, all classes of InDels and single nucleotide variants were down-sampled to 650,000 variants each.

A total of 854,409 *de novo* SNPs and indels were compiled from three different sources, lifted over to the hg38 assembly using the UCSC liftOver tool as required. First, we downloaded variants from http://denovo-db.gs.washington.edu/, including only samples from whole genome sequencing studies (Michaelson et al. 2012; Ramu et al. 2013; Genome of the Netherlands 2014; Besenbacher et al. 2015; Turner et al. 2016; Yuen et al. 2016; Jonsson et al. 2017; RK et al. 2017; Turner et al. 2017; Werling et al. 2018), which included a total of 404,238 variants from 4,560 samples. Additional samples, which were not already included in the denovo-db dataset, were downloaded from the MSSNG database (https://research.mss.ng/), version 2019/10/16, which added 2,243 samples and 215,044 *de novo* mutations. A third source of *de novo* variants came from(An et al. 2018) - 3,805 samples and 255,107 mutations.

### Circular Permutation

To obtain a genome-wide estimate of enrichment of overlap between genomic features (e.g. TFBSs and mutations), we performed circular permutations using the Bioconductor regioneR package in R (Gel et al. 2016). We used the permTest() function with parameters ntimes=10000, randomize.function=circularRandomizeRegions, evaluate.function=numOverlaps, genome=hg38_masked, alternative=“auto”, where hg38_masked = getBSgenome(“BSgenome.Hsapiens.UCSC.hg38.masked”). This test evaluates the number of overlaps observed between two sets of genomic features, given their order on the chromosome and the distance between features, i.e. taking their degree of clustering into account. At each iteration, all positions are shifted by the same randomly generated distance on conceptually “circularized” chromosomes, in effect “spinning” the position of features on each chromosome, while excluding masked regions (i.e. unmappable, repetitive and low-complexity segments). The statistical output is a z-score, which is defined as the distance between the expected number of overlaps (the distribution over 10,000 permutations) and the observed one, measured in terms of standard deviations. Shifted Z-score analysis can further indicate whether a given overlap pattern is caused by the precise locations of two feature sets or by broader, regional effects. One set of features is shifted in either direction from their original positions by a number of bases, so that the numbers of overlaps and the degree to which the z-score changes in response to the shift can be tested. A sharp peak of shifted z-scores around the zero-coordinate indicates a precise overlap between features, whereas a flat profile may indicate overlaps attributable to regional effects, as is seen for features that tend to co-occur in regions with similar base composition or gene density. For permutations involving SVs, we used the two breakpoints of each SV, and assessed the overlap of breakpoints with another feature of interest (i.e. ATAC-Seq sites), treating each breakpoint separately. Circular permutations in regioneR (Gel et al. 2016) were also used assess the mean distance between ATAC-Seq peaks and deletion breakpoints, for common and singleton variants separately.

### Simple permutations

Spermatogonial binding sites were randomly permuted, using the “bedtools shuffle” command (Quinlan and Hall 2010), with the parameter “-noOverlapping” and excluding assembly gaps with “-excl hg38.gap.bed” (downloaded from https://genome.ucsc.edu/cgi-bin/hgTables). In each of 10,000 simulations, we assessed the overlap of the permuted binding sites with short insertions and deletion breakpoints, respectively. This resulted in an expected distribution of overlap, given a random positioning of binding sites across the genome. This distribution was compared to the observed number of overlaps (using bedtools intersect), and the p- value was defined as the percentage of simulated overlaps that was larger than the observed overlap. This permutation framework does not take into account the spacing and clustered nature of binding sites, and does not allow for an assessment of the precision of overlap between features. In this study, we favor the more conservative measures of significance provided by the circular permutation strategy.

### Brain enhancer data

Active brain enhancers came from de la Torre-Ubieta et al. (2018). Specifically, we used the 37,050 brain enhancers which showed differential accessibility in the germinal zone versus the cortical plate, reflecting activity in the developing brain (de la Torre-Ubieta et al. 2018). Next, we identified brain enhancers that were also active during the male germline formation, i.e. overlapping the spermatogonial ATAC-Seq peaks. To correct for the variable size of the brain active enhancers, we took the midpoints of each enhancer plus/minus 500 bp on either side, and intersected these sites with the ATAC-Seq peaks using bedtools intersect (Quinlan and Hall 2010), thus classifying brain enhancers as spermatogonial “active” or “inactive”. Next, we intersected these two categories of brain enhancers with the DDD breakpoint and gnomAD insertion dataset, respectively, to further classify them as “disrupted” by a singleton variant or “intact”. An odds ratio was calculated as

OR = (A/(B - A))/(C/(D - C))

With confidence intervals

CI_lower = exp(log(OR) -1.96 * sqrt(1/A + 1/(B-A) + 1/C + 1/(D-C)))

CI_higher = exp(log(OR) +1.96 * sqrt(1/A + 1/(B-A) + 1/C + 1/(D-C)))

where:

A = Disrupted, sperm active

B = All sperm active

C = Disrupted, sperm inactive

D = All sperm inactive

To analyse the enrichment of short InDels and SNPs around TFBSs and brain enhancers, we only considered genomic regions with unique mappability in >= 95% of the region, using the bedmap option --bases-uniq-f (Neph et al. 2012) and the mappability file hg38_umap24 (Karimzadeh et al. 2018), converted to bedmap format.

### Random Forest Regression

To compare the effects of chromatin state on mutation rates, we performed random forest regression with 200 trees, modelling the outcome variables “singleton breakpoints” and “singleton insertions (5-20 bp)”, from the DDD and gnomAD V.3 respectively, within 5-kb wide genomic windows. Predictor variables included “spermatogonial TFBS count”, “ssDNA overlap” (from Pratto et al.(2014)), “recombination HS overlap” (from The 1000 Genomes Project et al. (2015)), “GC- content”, “Replication timing” (average of Wavelet-smooth signal in 1-kb bins of 15 encode tissues, downloaded from http://hgdownload.soe.ucsc.edu/goldenPath/hg19/encodeDCC/wgEncodeUwRepliSeq /), “Gene density”, “Mappability” (proportion of sites in each window with an umap24 score of 1), and the overlap with “LTRs”, “SINEs”, “LINEs” and “Simple Repeats” (downloaded from the Table Browser at https://genome.ucsc.edu/).

In a smaller model, we subsetted the dataset to only include 5-kb bins that also overlap active brain enhancers (de la Torre-Ubieta et al. 2018), then ran the random forest regression model to predict mutation rates within genomic regions that contain active brain enhancers.

### Motif discovery in singleton insertion sites

In order to find sequence motifs within the 5-20 bp singleton insertion sites from gnomAD V.3, without prior assumptions, we extracted the fasta sequence for insertions that fell within 10 bp of the top 10 disrupted motif families (motif families 992, 193, 796, 907, 579, 825, 984, 171, 991). We ran the meme.4.11 motif discovery algorithm (Bailey et al. 2009) with “-nmotifs 1” on the inserted sequences. This allowed us to compare the sequence motif of the disrupted TFBSs to any recurrent motif found within the inserted sequences.

### Control Motif sites

Using default search criteria, the FIMO algorithm (Grant et al. 2011) was run on the repeat masked hg38 genome sequence (hg38.fa.masked, downloaded from https://genome.ucsc.edu/ in March 2020), searching the whole genome for the 579 input Jaspar motifs and the 13-mer PRDM9 motif. As with active binding sites, motif matches belonging to the same motif family were merged and reported as a single motif match per family, and only regions with unique umap24 mappabilities for >= 95% of sites were kept; motifs that overlapped with spermatogonial ATAC-Seq peaks were excluded. Next, these “control” motif sites were down-sampled to 10,000 per motif family (using bedtools sample (Quinlan and Hall 2010)); circular permutations were performed to compare the observed to expected overlap of the control motif sites (plus/minus 10 bp) with the gnomAD singleton insertions of 5-20 bp.

The FIMO predicted control sites were also used to assess the degree of “clustering” of motifs at spermatogonia active sites. For this purpose, we intersected the FIMO motifs with a) spermatogonial ATAC-Seq sites and b) ENCODE Master regulatory sites downloaded from https://genome.ucsc.edu/ (DNaseI hypersensitivity derived from assays in 95 cell types). For each of the 167 motif families, we calculated the median distance (in basepairs) from a motif located within the active regulatory region to the nearest FIMO motif of the same type. Accordingly, the ratio of the median distance between motif sites (ENCODE/spermatogonia) was larger than one if motifs at spermatogonial sites were, on average, closer to each other than motifs near ENCODE sites, and we used this ratio as a measure of motif clustering. When correlating the IFE with the degree of motif clustering (Figure 6), we thus largely correct for base compositional biases near active sites (which impact mutation rates – Supplemental Fig. S9) as well as the effects of historical selection on the clustering of motifs near genes, i.e. shorter inter-motif distances in spermatogonia indicate that these sites have specifically high levels of motif density in spermatogonia, beyond the levels expected for binding sites in general.

### Data access

Raw sequencing data generated in this study have been submitted to the European Genome-phenome Archive (EGA) (accession number EGAS00001005366), and ATAC-Seq peak files are available at Edinburgh DataShare (https://doi.org/10.7488/ds/3053).

## Competing interest statement

The authors have no competing interests to declare.

## Acknowledgements

We thank all donors for their participation in genetic research. In particular, we thank the DDD families, study clinicians, research nurses and clinical scientists in the recruiting centres; the Genome Aggregation Database (http://gnomad.broadinstitute.org/), MSSNG (https://www.mss.ng/) and denovo-db (http://denovo-db.gs.washington.edu/denovo-db/) for making their data available. We are grateful to all of the families at the participating Simons Simplex Collection (SSC) sites, as well as the principal investigators (A. Beaudet, R. Bernier, J. Constantino, E. Cook, E. Fombonne, D. Geschwind, R. Goin-Kochel, E. Hanson, D. Grice, A. Klin, D. Ledbetter, C. Lord, C. Martin, D. Martin, R. Maxim, J. Miles, O. Ousley, K. Pelphrey, B. Peterson, J. Piggot, C. Saulnier, M. State, W. Stone, J. Sutcliffe, C. Walsh, Z. Warren, E. Wijsman). We appreciate obtaining access to genetic data on SFARI Base. Approved researchers can obtain the SSC population dataset described in this study by applying at https://base.sfari.org. This work was supported by MRC Human Genetics Unit core funding programme grants MC_UU_00007/11, MC_UU_00007/2 and MC_UU_00007/16. We thank Elisabeth Freyer for assistance with the FAC sorting and Wendy A. Bickmore for useful comments to the manuscript.

## Author Contributions

V.B.K. and C.A.S. conceived the project, interpreted the results and wrote the manuscript. M.S.T. designed the experiments and managed the acquisition of samples. L.T., Y.K., F.S. and M.M. performed the experiments, L.T. processed raw data. V.B.K. performed the analyses. D.D.D. and D.R.F. provided data. D.R.F. helped with the interpretation of the results and provided critical scientific inputs.

## Notes

### Competing Interest Statement

The authors have declared no competing interest.

