## Supplemental Figures for "Mutational bias in spermatogonia impacts the anatomy of regulatory sites in the human genome"

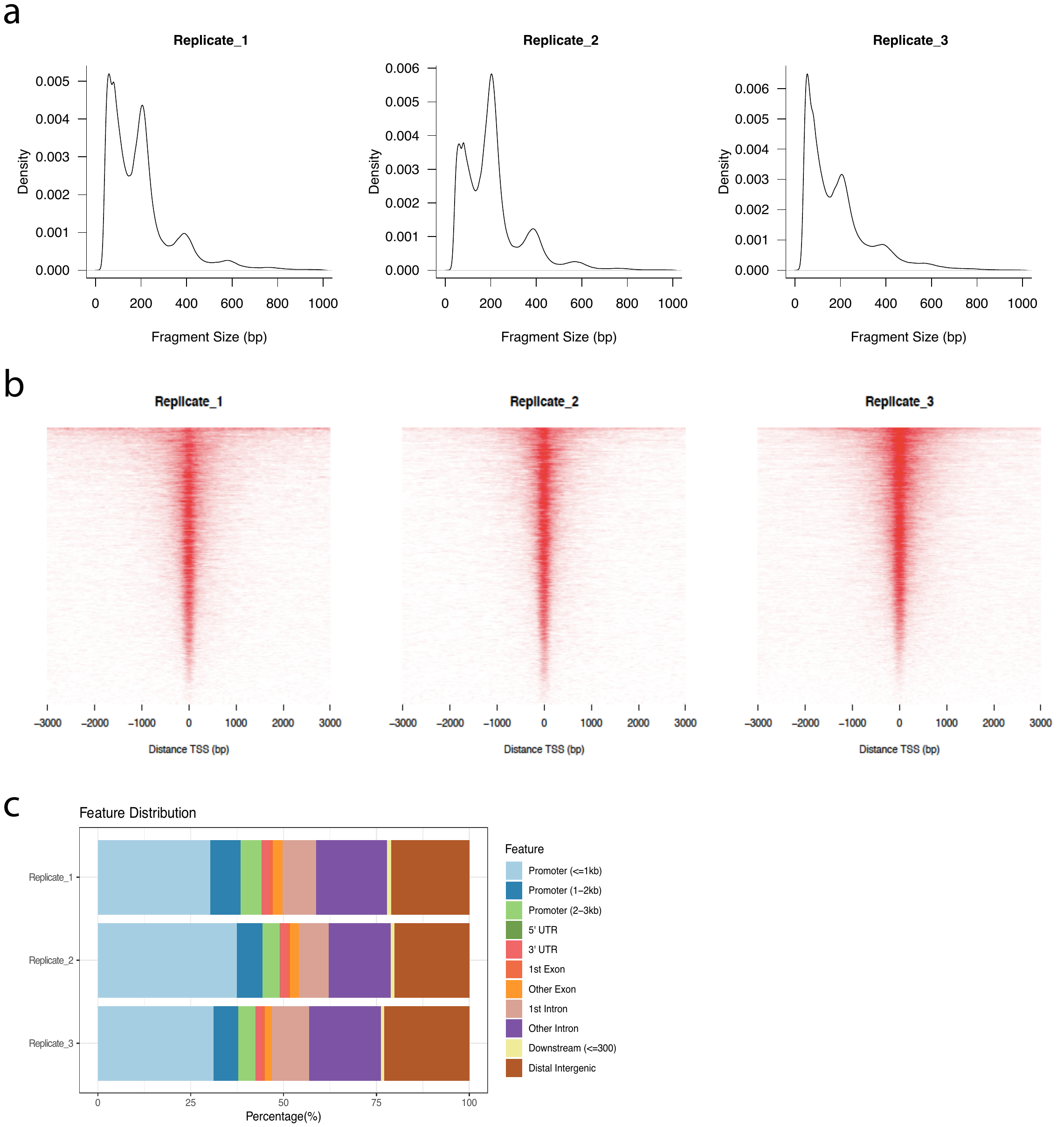

**Figure S1: Summary of spermatogonial ATAC-Seq data.** a) Fragment length distribution, showing the ~200bp periodicity around nucleosome-free regions; fragments <=100bp in length were chosen for downstream analysis and peak calling. b) Heatmap of ATAC-Seq peak density around transcription start sites (TSSs). c) Genomic feature annotation of ATAC-Seq peaks. b) and c) were produced using ChIPseeker, using the UCSC hg38 genome annotation.

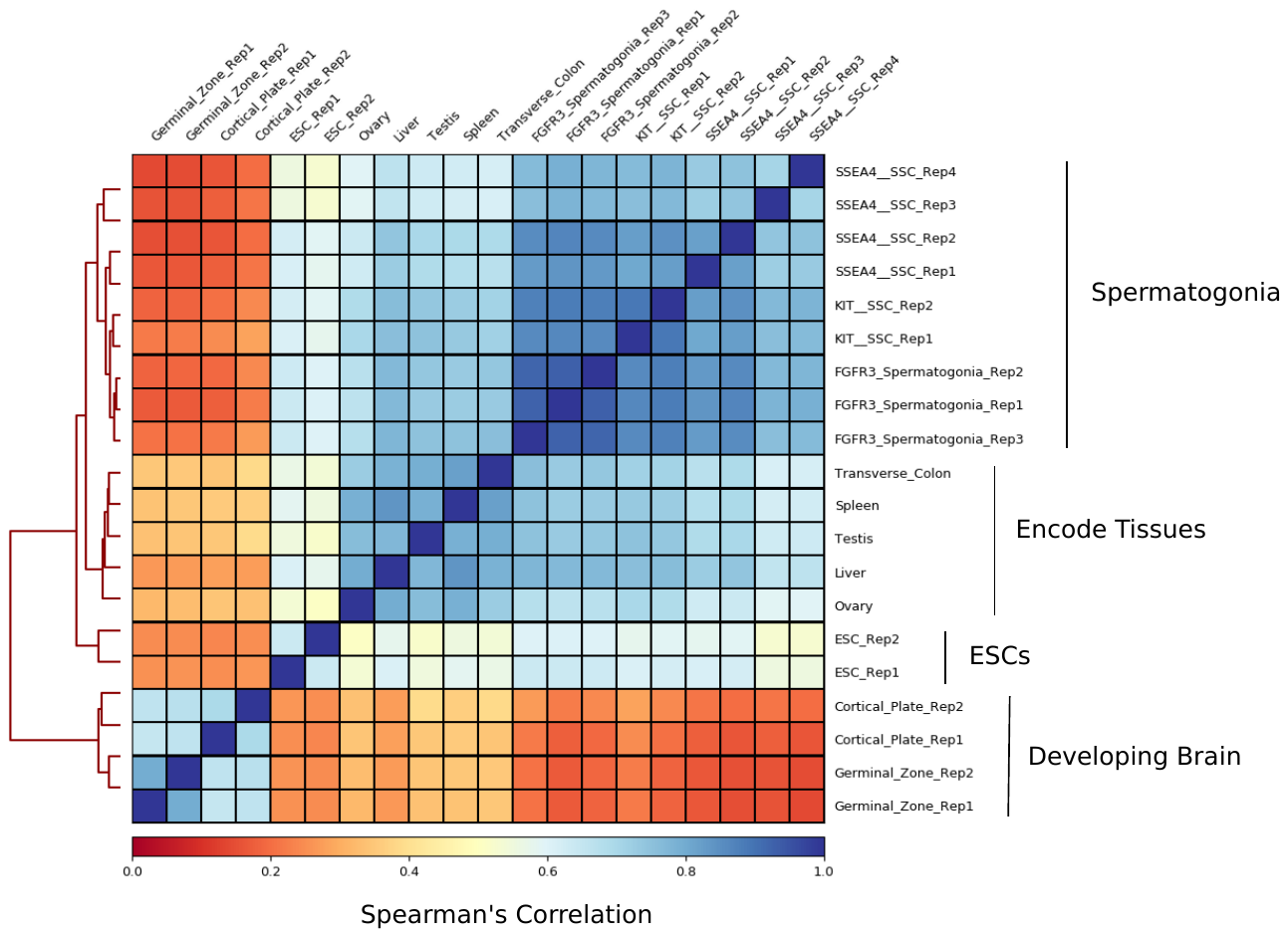

**Figure S2: Clustered heatmap of ATAC-Seq datasets.** Colours indicate the correlation coefficients between the genome-wide ATAC-Seq signals in

SSEA4+ spermatogonial stem cells (SSC) (Guo, Grow et al. 2017); KIT+ SSC (Guo, Grow et al. 2018); FGFR3+ spermatogonia (this dataset); Encode tissues (Encode Project Consortium 2012, Davis, Hitz et al. 2018); ESC cells (Guo, Grow et al. 2017); the germinal zone and cortical plate of the developing brain (de la Torre-Ubieta, Stein et al. 2018).

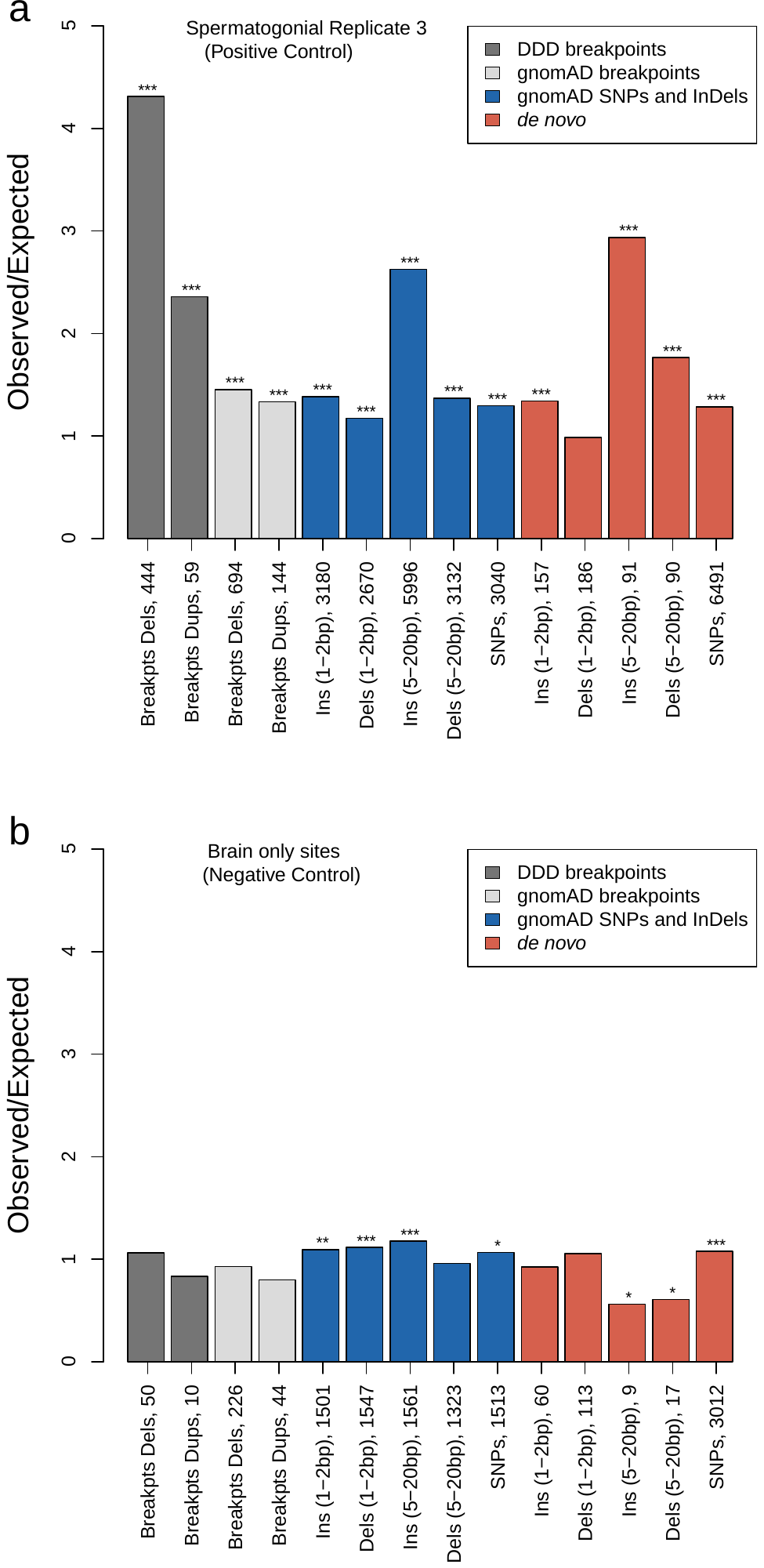

**Figure S3: The enrichment of short variants and SV breakpoints at spermatogonial binding sites, using positive and negative controls.** The Y axis shows the ratio of observed over expected variant counts at accessible sites, based on 10,000 circular permutations. Mutation categories with significant enrichment or depletion are indicated by asterisks (* = p < 0.05; ** = p < 0.01 *** = p < 0.001). The type of variant tested and the total number of observed variants overlapping TFBSs are indicated below each bar. a) Sites accessible in Replicate 3 (FGFR3+ spermatogonial cells). b) Sites that are accessible in the germinal zone and/or cortical plate (de la Torre-Ubieta et al. 2018), but not accessible in any of the spermatogonial samples (FGFR3-, SSEA4- or KIT-marked spermatogonia).

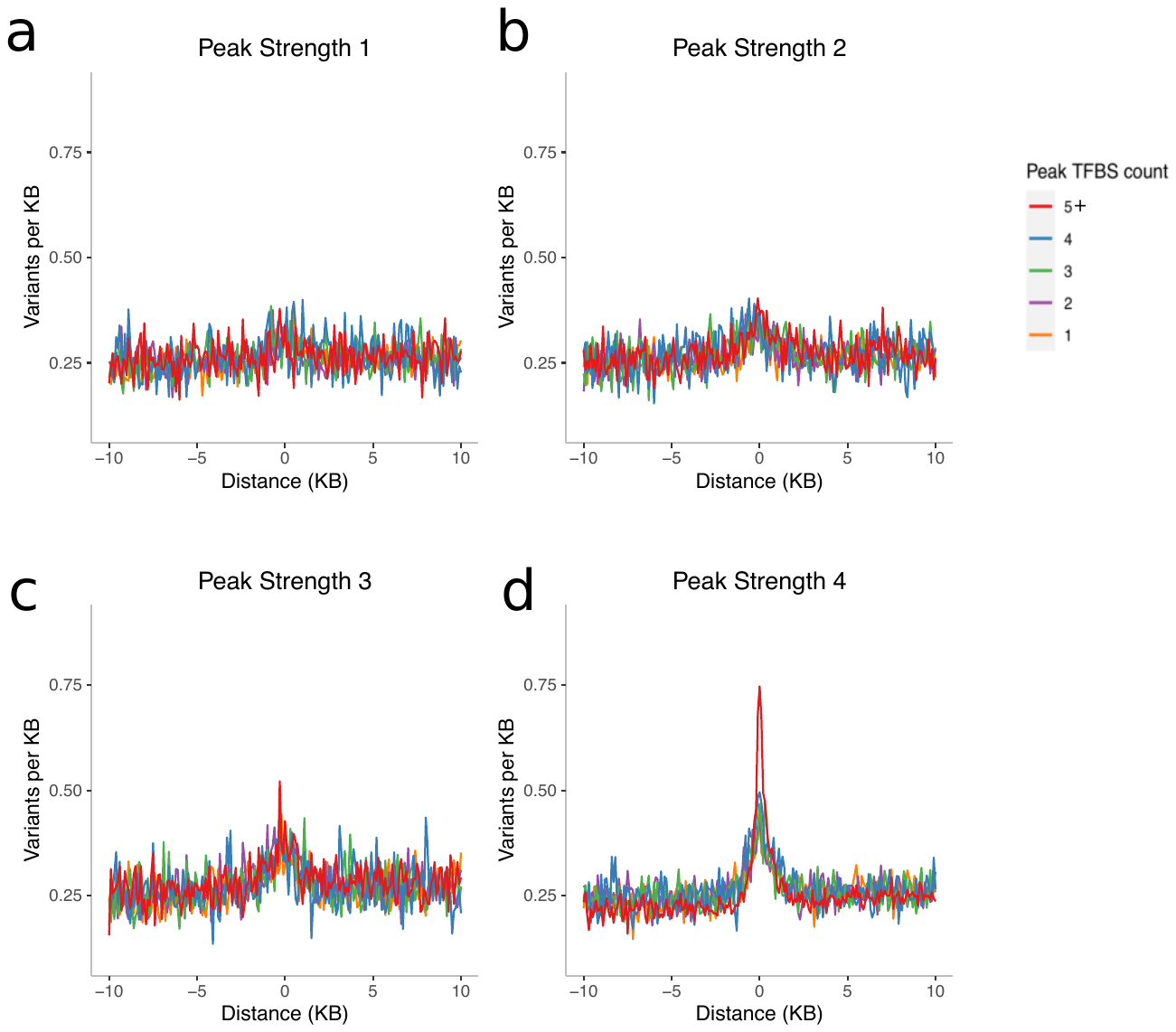
 **Figure S4: Short (5-20 bp) insertion rates, stratified by peak strength and the number of TFBSs within peaks. a-d)** Spermatogonial ATAC-Seq peaks were divided into four equal sized groups, based on their quartile of peak score (after removal of outliers, i.e. the top 1% of peak scores): Peak Strength 1, 2, 3 and 4. Next, the peaks were classified by the number of putative binding sites that they contained, ranging from 1 TFBS to “5 or more” TFBSs. The X axis shows the distance from the centre of the ATAC-Seq peak, and the Y axis shows the average rate of singleton insertions ( 5-20bp ) in the gnomAD dataset, for each category of ATAC-Seq peak.

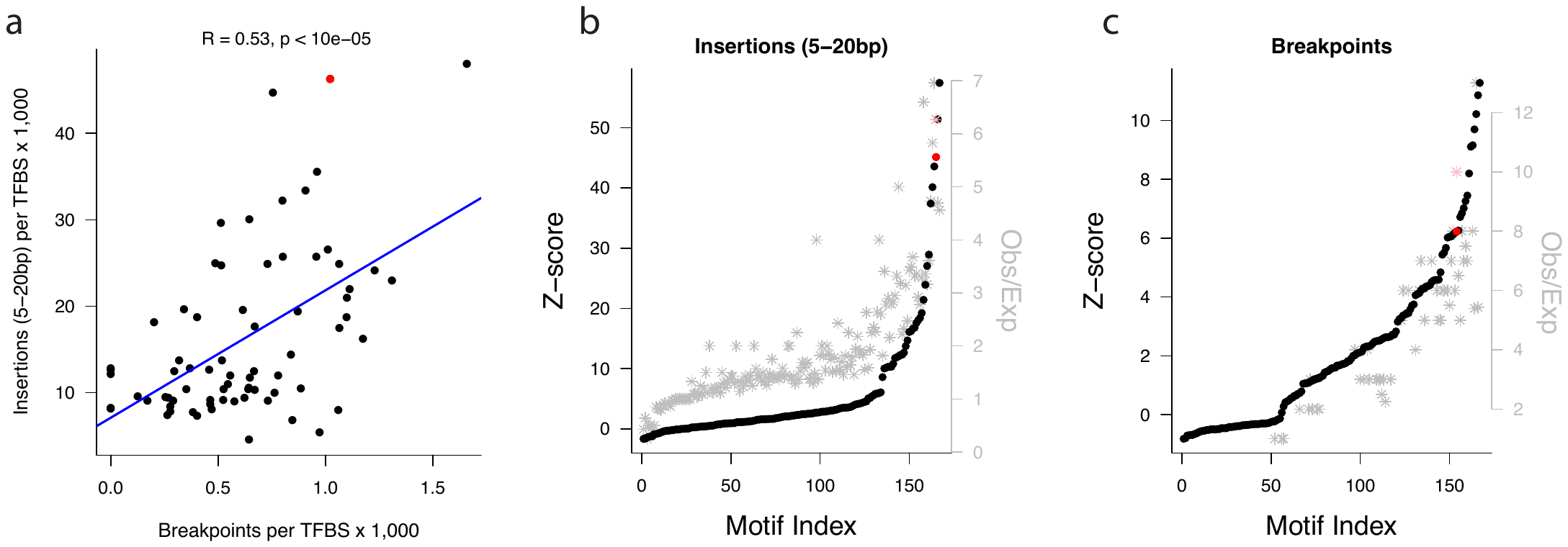

**Figure S5:** **Short insertions (5-20 bp) and deletion breakpoints are correlated.** **a)** Rates of singleton gnomAD insertions and DDD deletion breakpoints, for 72 motif families with at least 3,000 predicted binding sites in spermatogonia; each data point represents one motif family. PRDM9 is highlighted in red. **b, c,** Circular permutation Z-scores (left axis, black dots) and the enrichment of variants for each motif family (the ratio of observed over expected numbers of variants; right axis, grey stars). PRDM9 is highlighted in red and pink, respectively. Motifs are ranked by ranked by Z-score (Motif Index).

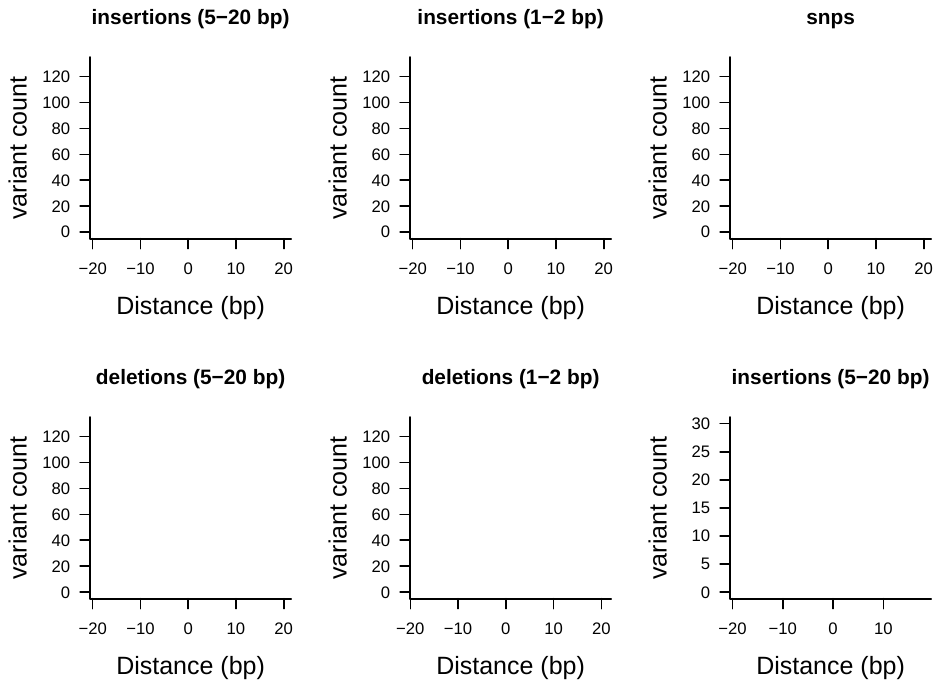

**Figure S6: SV deletion breakpoints often precisely overlap InDels (5-20bp).** The x-axis is centered around deletion breakpoints of the gnomAD structural variant dataset. Each data point indicates the aggregate number of gnomAD short variants (SNPs and InDels, respectively) observed at a given distance from the breakpoint, at basepair resolution. **Red**: singleton short variants *versus* singleton deletion breakpoints. All singleton short variants were down-sampled to a total of 650,000 variants each, making the Y axes comparable. **Blue**: High frequency short insertions (5-20bp) (*p* >= 5%) *versus* high frequency deletion breakpoints (*p* >= 5%).

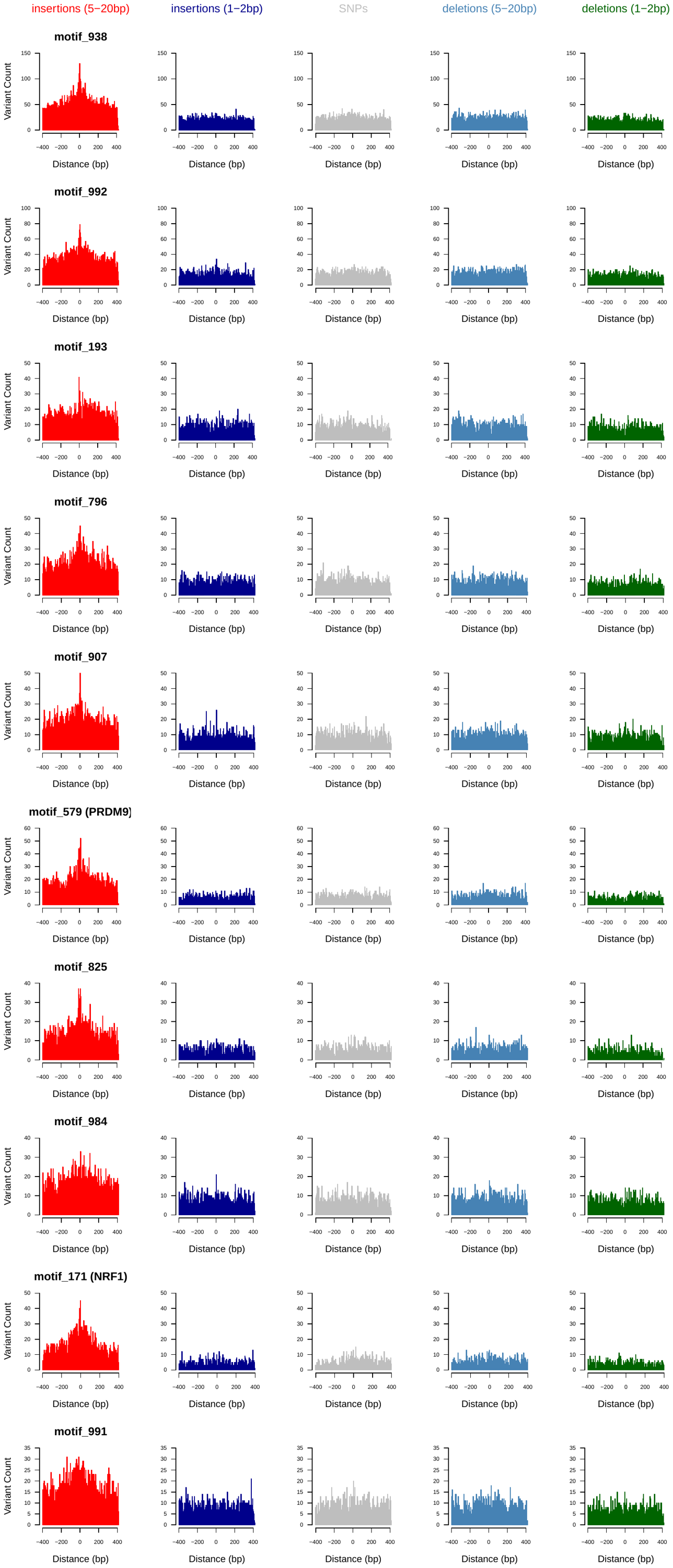
­­­

**Figure S7**: **Rate of singleton InDels and snps in the gnomAD dataset near active spermatogonial motif sites.** Insertions, Deletions and SNPs were down-sampled to a total of 650,000 variants each, making the Y axes comparable; individual bins are 5bp in size. The ten motifs shown had the highest overall number of observed insertions of 5-20 bp (Supplementary Table 4). Only regions around TFBSs with >=95% unique mappability (umap24 scores) were included.

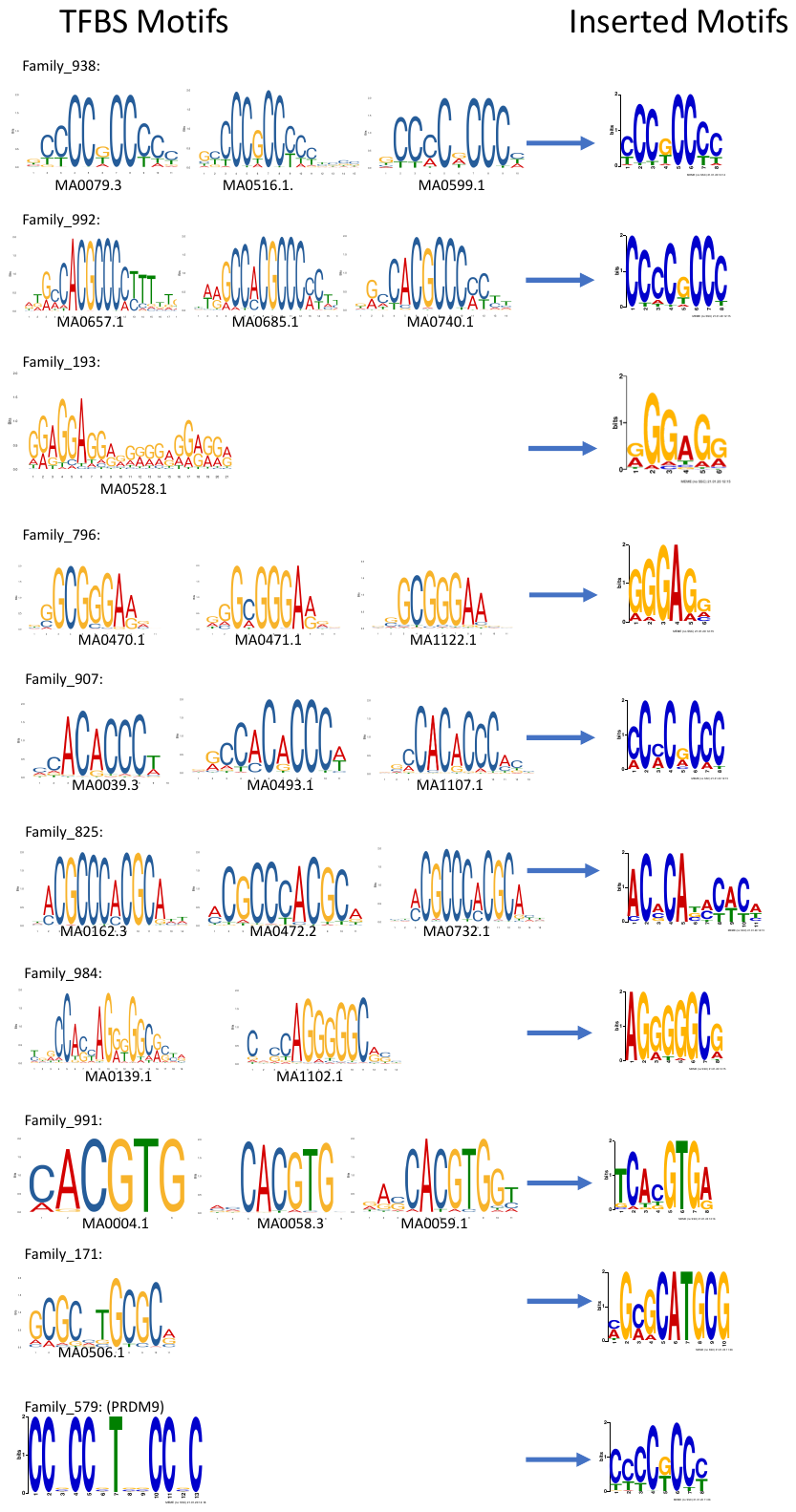

**Figure S8: Insertions often cause duplications of the binding motif.** Jaspar database sequence motifs in the footprints of spermatogonial ATAC-Seq peaks (left) and the motifs identified by meme in the singleton insertions (5-20bp) at these sites (right). Shown are the ten motif families with the overall highest number of insertions. A maximum of three Jaspar motifs per motif family are shown on the left.

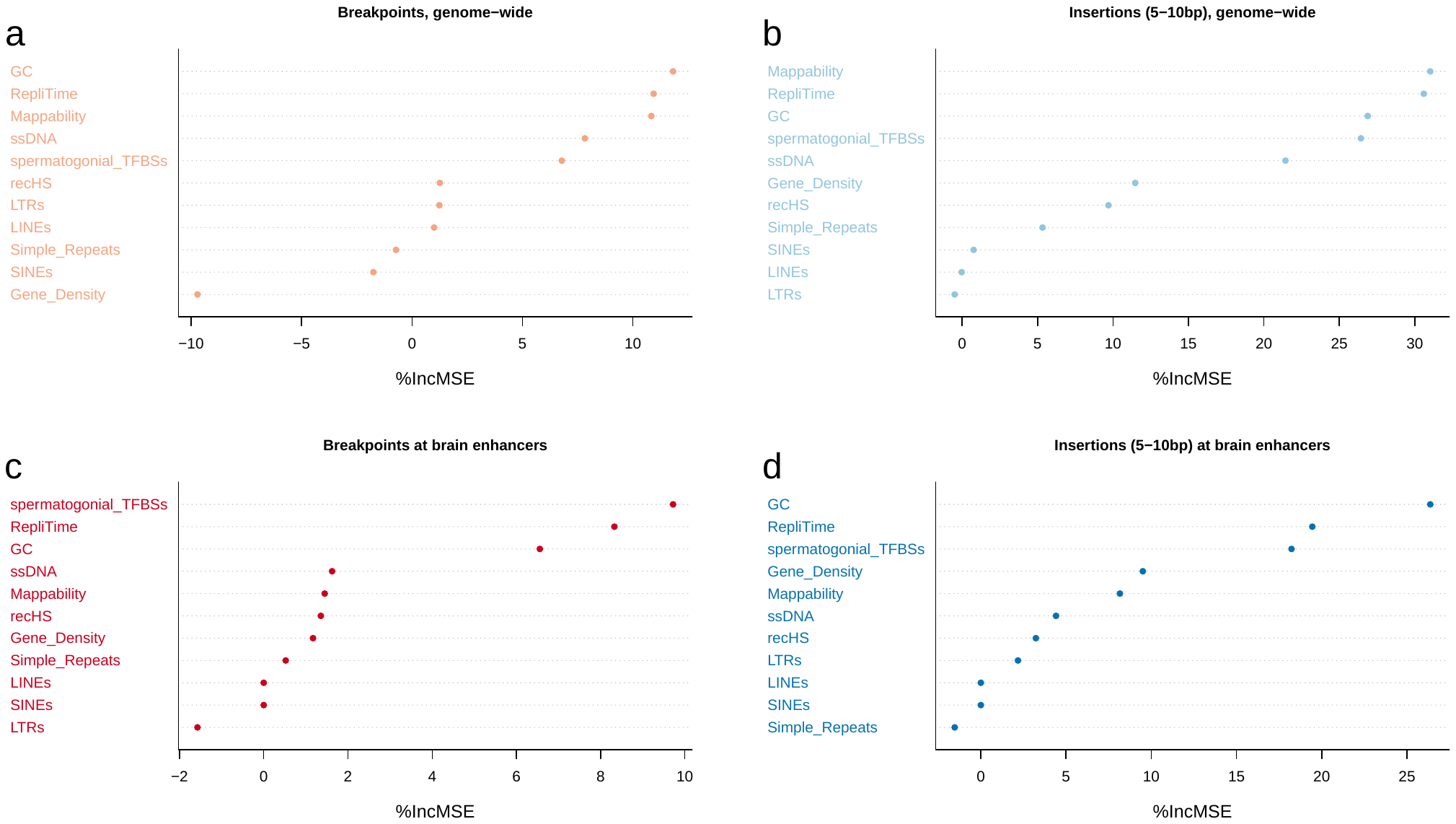

**Figure S9: Random Forest analysis of insertion (5-20 bp) and deletion breakpoint rates.** The importance of predictor variables, measured as the % increase in mean square error when the variable is removed from the model (%IncMSE), is plotted for four different random forest regression models. **a, b,** modelling mutation rates genome-wide (in 5 KB-wide bins). **c, d** modelling mutation rates in 5 KB-wide bins which also overlap brain active enhancers as defined in the main text.

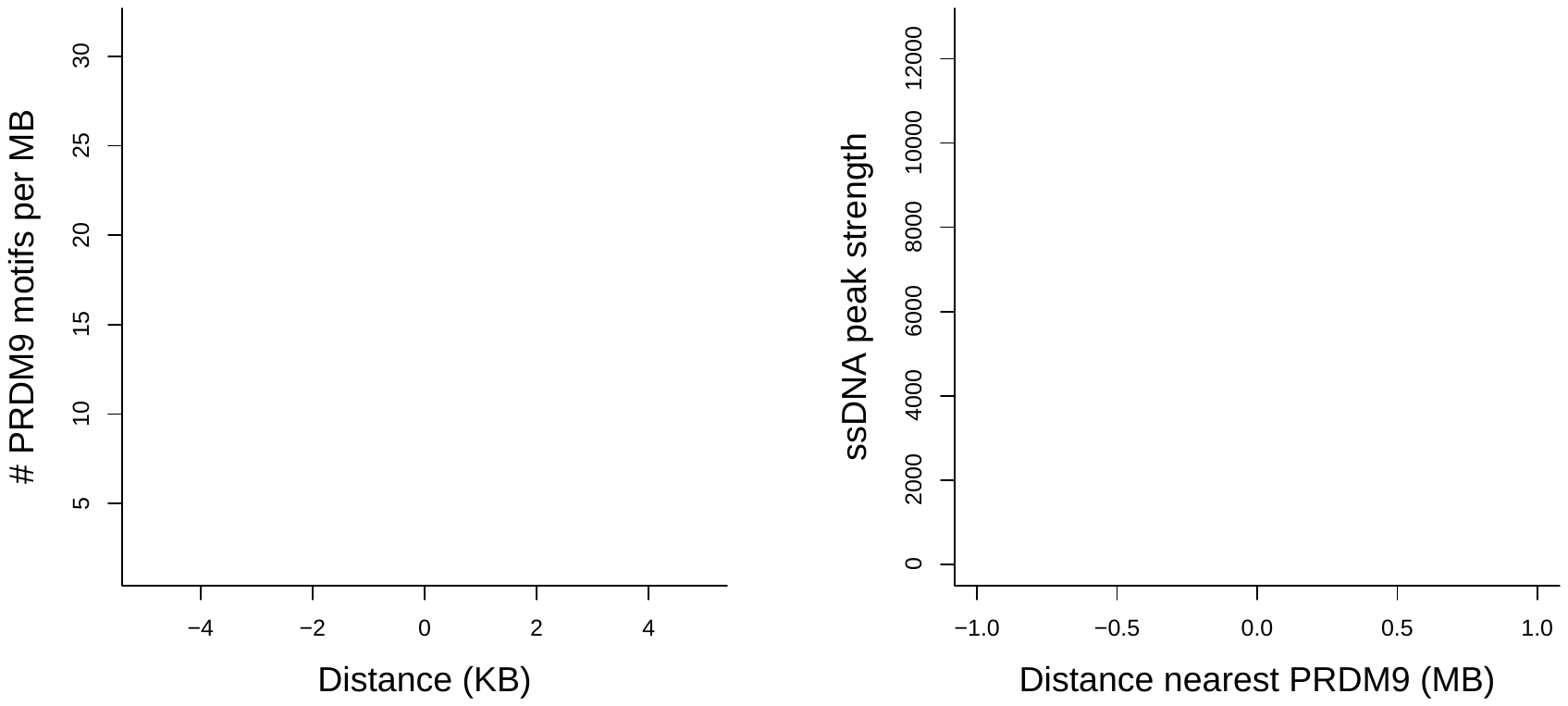

**Figure S10: PRDM9 motifs within ATAC-Seq footprints are enriched at testis-derived ssDNA sites.** **a)** The frequency of PRDM9-binding sites, centred around ssDNA sites. **b)** The ssDNA peak strength *versus* the distance to the nearest PRDM9-binding site. ssDNA data came from Pratto, Brick et al. (2014) and PRDM9-binding sites from the spermatogonial footprinting analysis.
